# RNase H enables efficient repair of R-loop induced DNA damage

**DOI:** 10.1101/068742

**Authors:** Jeremy D. Amon, Douglas Koshland

**Affiliations:** Department of Molecular and Cell Biology, University of California, Berkeley, CA 94720

## Abstract

R-loops, three-stranded structures that form when transcripts hybridize to chromosomal DNA, are potent agents of genome instability. This instability has been explained by the ability of R-loops to induce DNA damage. Here, we show that persistent R-loops also compromise DNA repair. Depleting endogenous RNase H activity impairs R-loop removal in budding yeast, causing DNA damage that occurs preferentially in the repetitive ribosomal DNA locus (rDNA). We analyzed the repair kinetics of this damage and identified mutants that modulate repair. Our results indicate that persistent R-loops in the rDNA induce damage that is slowly repaired by break-induced replication (BIR). Furthermore, R-loop induced BIR at the rDNA leads to lethal repair intermediates when RNA polymerase I elongation is compromised. We present a model to explain how removal of R-loops by RNase H is critical in ensuring the efficient repair of R-loop induced DNA damage by pathways other than BIR.

## Introduction

R-loops are structures that form when RNA invades double stranded DNA and hybridizes to complementary genomic sequences [1]. R-loops can form spontaneously across many genomic loci, but the activity of two endogenous RNases H prevents their accumulation and persistence [2]. RNase H1 and H2 are highly conserved ribonucleases with the ability to degrade the RNA moiety of a DNA:RNA hybrid. Disrupting the activity of the two enzymes (*rnh1*Δ *rnh201*Δ in yeast) has been a useful tool for increasing the persistence of DNA:RNA hybrids and studying the effects of hybrid-induced instability. Indeed, efforts to map R-loops genome-wide have shown that in the absence of RNase H activity, the levels of hybrids formed at spontaneous loci increase dramatically [3, 4]. This increase in hybrids is associated with increased rates of genome instability that include loss of heterozygosity (LOH) events, loss of entire chromosomes, and recombination at the ribosomal locus [4,5]. The RNases H have therefore been implicated as important protectors of genome stability.

The ribosomal locus (rDNA) appears to be particularly prone to R-loops. Approximately 60% of all transcription in the cell is devoted to producing ribosomal RNA from about 150 repeated units located in a clustered region on chromosome XII [6]. These repeats, at 9.1 kb each, make up about 10% of the yeast genome. Accordingly, almost 50% of all R-loops map to the rDNA [3]. R-loops found at the rDNA are associated with increased rates of recombination [4,7], RNA polymerase pileups [8], and stalled replication forks [9].

A growing body of evidence has attributed various biological roles to R-loops, including modifying gene expression [10,11], terminating transcription [12,13], driving sequence mutation [14], and inducing changes in genome structure [15,16]. However, the mechanisms of R-loop induced genome instability remain elusive. Most studies on the mechanisms of hybrid-induced instability have been “damage-centric,” investigating how R-loops are converted to mutations, single stranded nicks, and double stranded breaks (DSBs) [17]. Current models focus on the involvement of active replication forks that stall or collapse upon encountering the aberrant structure. While this remains an area of active research, we note that any instability event is the result of a complex interplay between the initial damage event and the repair processes that follow. Phenotypes that involve the loss of genetic information (terminal deletions, certain LOH events) imply both that damage occured and that repair processes failed to accurately maintain the genome. Few studies have investigated how R-loop induced damage is repaired, and it remains possible that defects in repair contribute to instability. This possibility raises several questions. First, do genomic changes induced by R-loops reflect increases in damage events, failures of repair, or both? Second, are specific pathways involved in the repair of R-loop induced damage, and if so, what are they?

To begin to answer these questions, we turned to the Rad52-GFP foci system in yeast. Rad52 is required in almost all homologous recombination pathways, and in yeast forms bright foci upon induction of DNA damage [18]. Most foci appear in the S/G2-M phases of the cell cycle and have a moderate rate of repair – almost all spontaneous Rad52-GFP foci dissipate within 40 minutes [19]. Consistent with phenotypes of increased genomic instability, *rnh1*Δ *rnh201*Δ mutants display an increase in Rad52-GFP foci. A large fraction of these foci appear to co-localize with the nucleolus and form in a window between late S and mid-M [9]. Here, by monitoring the persistence of Rad52 foci across the cell cycle in RNase H mutants, we implicate DNA:RNA hybrids in the disruption of DNA damage repair. We show that topoisomerase I works at the rDNA to prevent these disruptions from becoming lethal events. Furthermore, we identify a new role for the RNases H in preventing break-induced replication (BIR) from repairing R-loop induced DNA damage.

## Results

### The presence of either RNase H1 or H2 prevents the accumulation of DNA damage in G2-M

To better understand the mechanisms by which DNA:RNA hybrids contribute to genome instability, we began by characterizing DNA damage in exponentially dividing wild-type, *rnh1*Δ, *rnh201*Δ, and *rnh1*Δ *rnh201*Δ budding yeast cells. Using Rad52-GFP foci as a marker for DNA damage, we observed that 27% of *rnh1*Δ *rnh201*Δ cells had foci, a ten-fold increase over wild type, *rnh1*Δ, and *rnh201*Δ cells (Figure 1A). Consistent with the notion that persistent DNA damage uniquely affects the double mutants, the growth of the double mutant, but not either of the single mutants, was dramatically impaired by the deletion of *RAD52* (Figure 1B). Previous characterization of the double mutant also reported elevated foci and Rad52-dependent growth [9,20]. Thus, by measures of Rad52-GFP foci and Rad52-dependent growth, cells lacking RNase H1 and H2 have a larger fraction of persistent R-loop induced damage than wild-type cells or cells lacking only one of the RNases H. This persistent damage could have arisen from increased R-loop induced damage and/or an inability to efficiently repair that damage.

**Figure 1.**
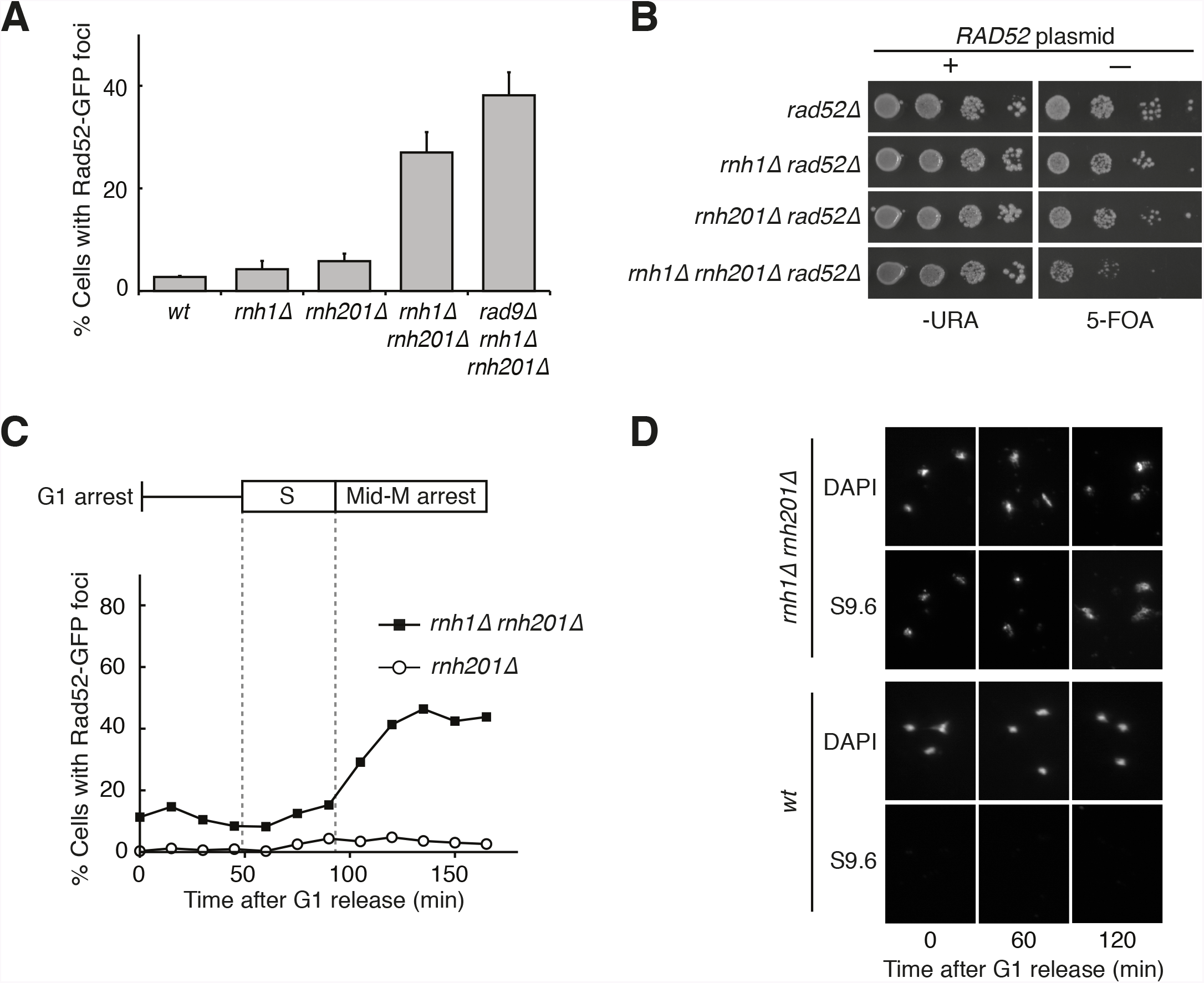
Cells lacking both RNases H accumulate DNA damage in G2-M. (A) Assessment of Rad52-GFP in RNase H mutants. Asynchronously dividing cells were scored for the presence of one or more Rad52-GFP focus. Bars represent mean +/- standard deviation (*n=3*; 300 cells scored per replicate). (B) Assessment of Rad52 requirement in RNase H mutants. Cells carrying a plasmid expressing RAD52 and URA3 were plated onto media lacking uracil (-URA, selects for plasmid) or media containing 5-floroorotic acid (5-FOA, selects for plasmid loss). 10-fold serial dilutions are shown. (C) Cell cycle profile of Rad52-GFP foci in RNase H mutants. Synchronously dividing cells were scored for the presence of Rad52-GFP foci. Cells arrested in G1 using alpha factor were washed and released into nocodazole. Samples were taken at 15-minute intervals and 300 cells per time point were scored for Rad52-GFP foci. Cell cycle phase is determined by flow cytometry (Supplemental Figure 2A). (D) Cell cycle profile of DNA:RNA hybrids in RNase H mutants. Shown are representative images of chromosome spreads of rnh1Δ rnh201Δ and wild-type cells. Spreads are stained for DNA content (DAPI) or immunostained for DNA:RNA hybrids using the S9.6 antibody and a fluorescent-conjugated secondary.

To further characterize the DNA damage response in *rnh1*Δ *rnh201*Δ cells, we asked whether this damage accumulated within a specific window of the cell cycle. We arrested *rnh1*Δ *rnh201*Δ cells in G1 using the mating pheromone alpha factor and released them into nocodazole, allowing them to proceed synchronously through the cell cycle until they arrested in mid-M phase at the spindle checkpoint (Figure 1C, Supplemental Figure 1A). During this cell cycle progression, aliquots of cells were removed and fixed to assess Rad52-GFP foci accumulation. Cell cycle stage was determined by measuring DNA content using flow cytometry (Supplemental Figure 1A). The fraction of cells with Rad52-GFP foci remained around 10 to 15 percent through S-phase. Additional foci appeared at the S/G2-M boundary and accumulated to around 50 percent, as reported previously. The failure to observe accumulating foci early in the cell cycle was not a limitation of the system, as an identical analysis of a single cell cycle of *sin3Δ* cells, which also accumulate hybrids, revealed an increase in focus formation during S-phase (Supplemental Figure 2A, 2B). The increase in foci in *rnh1*Δ *rnh201*Δ cells did not appear to be due to a cell-cycle dependent increase in hybrid formation, as cytological staining revealed similar levels of R-loops in cells staged in G1, S and M (Figure 1D). Therefore, the increase in damage during the S/G2-M window in *rnh1*Δ *rnh201*Δ cells is likely because hybrids were either more efficiently converted to damage or the repair of hybrid-induced damage became impaired.

**Figure 2.**
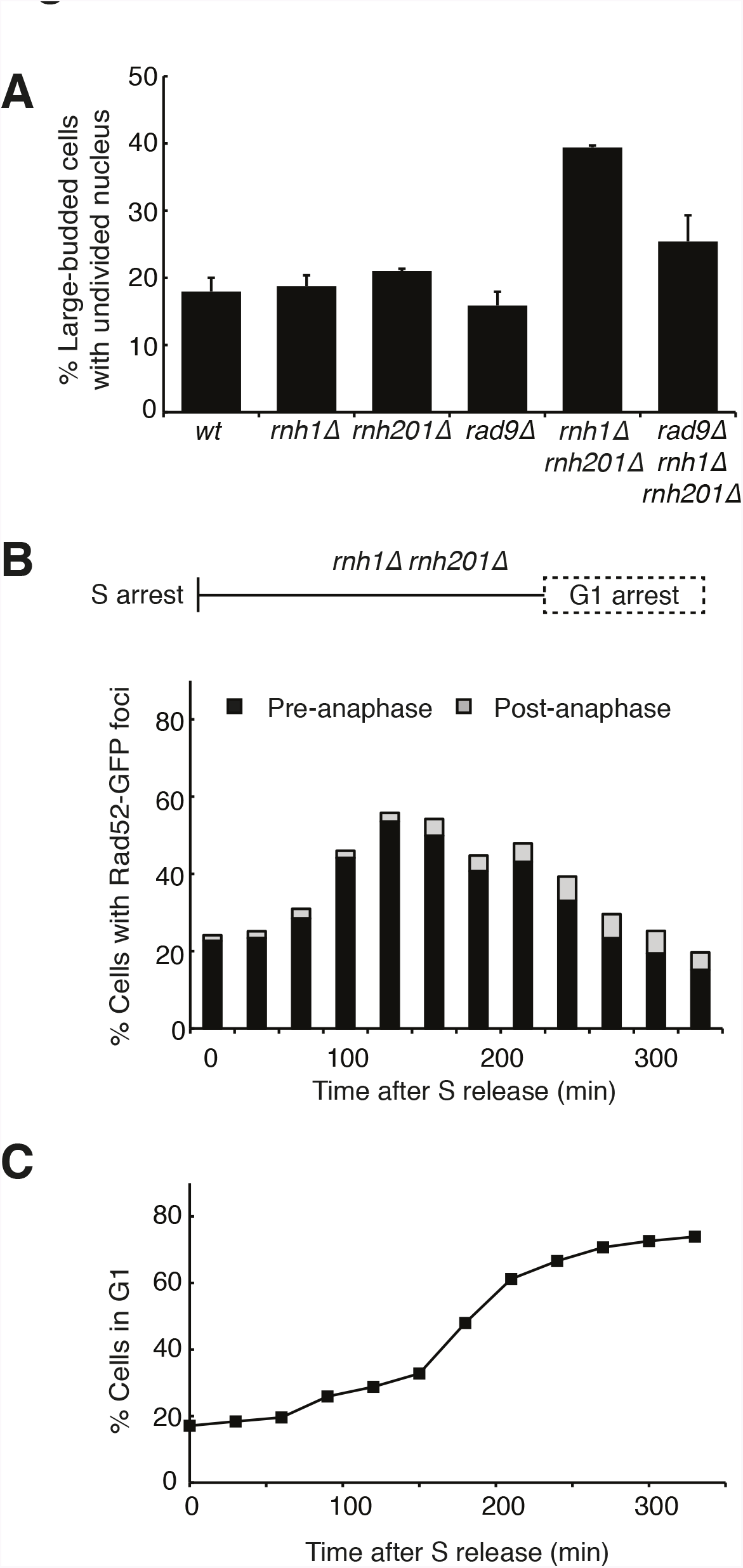
Cells with hybrid-induced DNA damage are slow to repair. (A) Assessment of cell-cycle delay in RNase H mutants. Asynchronously dividing cells were scored on the basis of their bud size and nuclear morphology. The percentage of cells with large buds and an undivided nucleus (single DAPI mass) are shown. Bars represent mean +/- standard deviation (*n=3*, 100 cells scored per replicate) (B) Cell cycle profile of Rad52-GFP foci in dividing cells. rnh1Δ rnh201Δ cells were arrested in S-phase using hydroxyurea, washed, and released into alpha factor. Samples were taken at 30-minute intervals and 300 cells per time point were scored for Rad52-GFP foci. If a cell had a Rad52-GFP focus, it was further scored for cell cycle phase. Cells with undivided nuclei (single DAPI mass) are labeled “pre-anaphase,” while those that had undergone nuclear division (two DAPI masses or G1 arrested) are labeled “post anaphase.” (C) Cell cycle stage of dividing rnh1Δ rnh201Δ cells. Cells from (B) were subjected to flow cytometry (Supplemental Figure 2c) and quantified. The percentage of cells with 1C DNA content is shown.

The presence of DNA damage such as DSBs leads to a Rad9-dependent cell-cycle checkpoint that delays entry into anaphase [21]. We found that the fraction of cycling cells in G2-M, defined as a large-budded morphology with an undivided nucleus, was two-fold higher in *rnh1*Δ *rnh201*Δ cells than wild type or either RNase H single mutant. This fraction was reduced by deletion of *RAD9* (Figure 2A). Deletion of *RAD9* did not decrease the level of Rad52-GFP foci in *rnh1*Δ *rnh201*Δ cells, indicating that focus formation is not dependent on the checkpoint (Figure 1A). To assess the kinetics of foci persistence in *rnh1*Δ *rnh201*Δ cells, we arrested cultures in S-phase using hydroxyurea and released them into alpha factor, allowing them to proceed through M-phase and arrest in the following G1 (Figure 2B and Supplemental Figure 1B). After the expected increase of Rad52-GFP foci upon the completion of S-phase, we observed a gradual disappearance of foci. Throughout the time-course, the vast majority of cells that retained foci were arrested pre-anaphase, indicating that most cells delayed progression into anaphase until the damage was repaired (Figure 2B). For example, after 330 minutes, the bulk of cells had reached G1 (Figure 2C) and the fraction of cells with Rad52-GFP foci had dropped to 20 percent. Of the cells that retained foci, 77 percent remained arrested in G2-M before anaphase. The slow disappearance of foci and progression into anaphase raised the possibility that hybrid-induced damage might be difficult to repair in a subset of the double mutant cells.

### Depletion of topoisomerase-1 exacerbates DNA damage phenotypes in the absence of the RNases H

To improve our ability to interrogate the unusual DNA damage in *rnh1*Δ *rnh201*Δcells, we sought to strengthen the damage phenotype. A number of observations suggested that alleles of *TOP1*, which encodes the major topoisomerase I in yeast, might be good candidates for doing so. Top1 is thought to clear R-loops and stalled RNA polymerase I (RNA pol I) complexes at the ribosomal locus by resolving supercoiling [8,22,23]. A potential synergistic relationship between Top1 and the RNases H came from the observation that while cells with either the *top1Δ* mutation or the *rnh1*Δ *rnh201*Δ mutations are viable, the *top1*Δ *rnh1*Δ *rnh201*Δ mutant is inviable [8]. Furthermore, treatment of *rnh1*Δ *rnh201*Δ cells with the Top1 inhibitor camptothecin led to increased Rad52-GFP foci that co-localized with the nucleolus [9]. Encouraged by these results, we used the auxin-inducible degron (AID) system to create a conditional *TOP1-AID* allele in wild type, the two single RNase H mutants, and the double RNase H mutant. We then reassessed viability and DNA damage phenotypes.

Consistent with published results,*rnh1*Δ *rnh201*Δ *TOP1-AID* cells fail to grow when treated with auxin (Figure 3A). In contrast, *TOP1-AID*, *rnh1*Δ *TOP1-AID*, and *rnh201*Δ *TOP1-AID* mutants grew well. Thus, the synergistic lethality occurred only when both RNases H and Top1 were inactivated. Similarly, when exponential cultures of these strains were treated with auxin for four hours, Rad52-GFP foci did not increase in *TOP1-AID*, *rnh1*Δ *TOP1-AID* or *rnh201*Δ *TOP1-AID* mutants (Figure 3B). However, foci nearly doubled in the *rnh1*Δ *rnh201*Δ *TOP1-AID* cells compared to an untreated control, such that a large majority of *rnh1*Δ *rnh201*Δ *TOP1-AID* cells (85%) had foci. Furthermore, after four hours of treatment with auxin, over 98 percent of *rnh1*Δ *rnh201*Δ *TOP1-AID* cells were arrested pre-anaphase at the G2-M checkpoint (Figures 3C and 3D). This arrest reflected an exacerbation of the cell cycle delay observed in the *rnh1*Δ *rnh201*Δ strain and in *rnh1*Δ *rnh201*Δ *TOP1-AID* cells left untreated with auxin (Figures 2A and 3C). As with *rnh1*Δ *rnh201*Δ cells, the cell-cycle arrest of the *rnh1*Δ *rnh201*Δ *TOP1-AID* was Rad9 dependent; deletion of *RAD9* resulted in cells that proceeded into the following G1. Importantly, deletion of *RAD9* did not restore viability to *rnh1*Δ *rnh201*Δ *TOP1-AID* cells treated with auxin. This result suggests that the inviability of the triple mutant was not simply due to the constitutive activation of the checkpoint but rather to the presence of irreparable damage.

**Figure 3.**
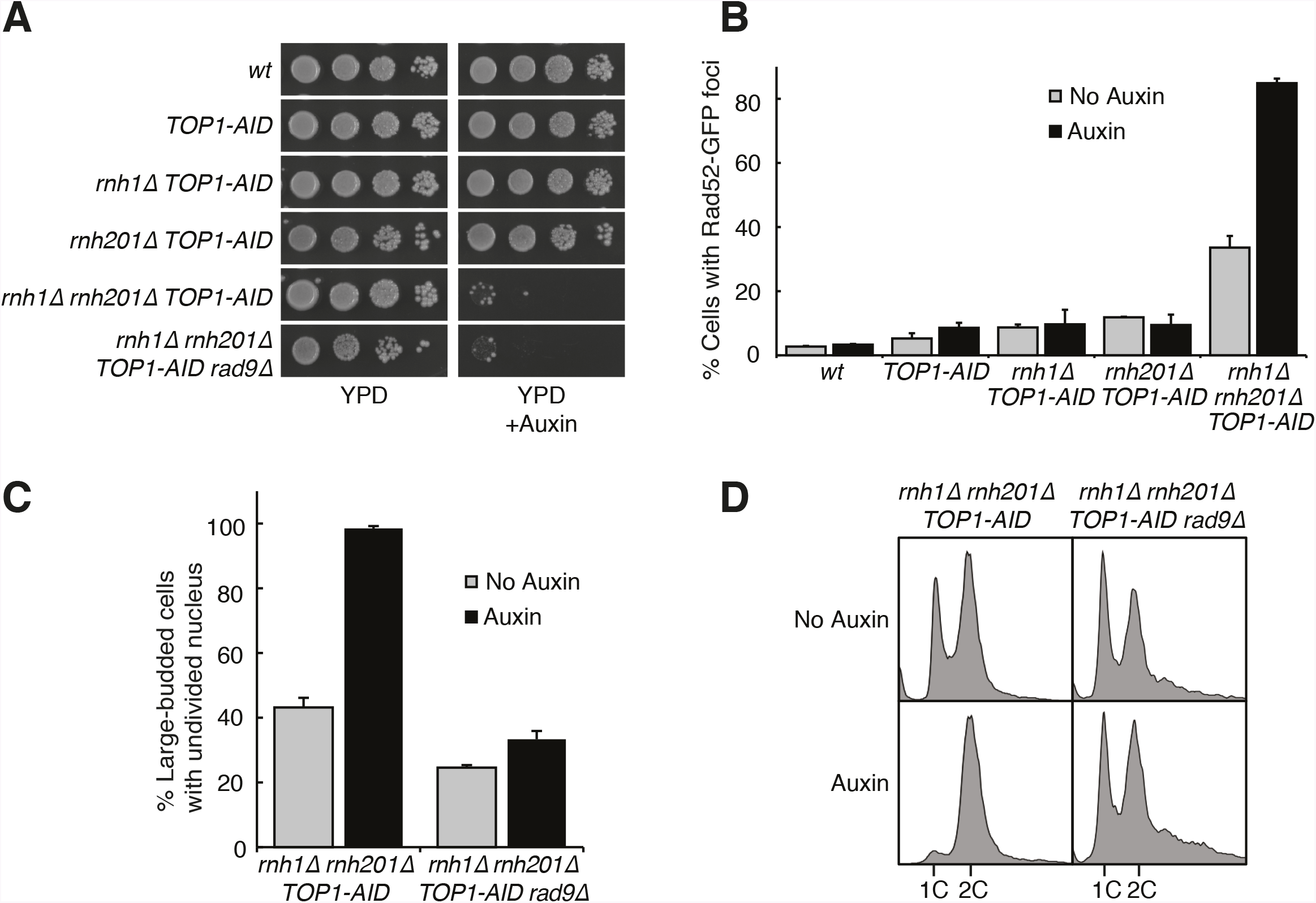
Depleting topoisomerase I exacerbates rnh1Δ rnh201Δ phenotypes. (A) Assessment of Top1 depletion on viability of RNase H mutants. 10-fold serial dilutions of saturated cultures were plated onto rich media (YPD) or media containing auxin (YPD +Auxin). (B) Assessment of Top1 depletion on Rad52-GFP foci in RNase H mutants. Cultures were grown at 23 degrees and treated with auxin for four hours. Cells were then scored for presence of Rad52-GFP foci. Bars represent mean +/- standard deviation (*n=3*, 300 cells scored per replicate). (C) Depleting Top1 leads to robust Rad9-dependent cell cycle arrest. Logarithmically dividing cells were treated with auxin for four hours then scored for bud size and nuclear morphology. The percentage of cells with large buds and undivided nuclei (single DAPI mass) is shown. Bars represent mean +/- standard deviation (*n=3*, 100 cells scored per replicate). (D) Cells from (C) were subjected to flow cytometry.

A striking feature of the Rad52-GFP foci in the *rnh1*Δ *rnh201*Δ double mutant was that they accumulated in a window that began at the boundary between S and G2-M. Therefore, we tested whether the enhanced focus formation in the *rnh1*Δ *rnh201*Δ *TOP1-AID* cells also occurred in this window (Figure 4A). A culture of the *rnh1*Δ *rnh201*Δ *TOP1-AID* triple mutant was arrested in G1 with alpha factor and treated with auxin to deplete Top1-AID (Supplemental Figures 3A and 3C). Cells were released from G1 into media containing auxin and nocodazole, to perpetuate Top1-AID depletion and induce subsequent arrest in mid-M (Supplemental Figure 3A). Aliquots were removed as cells progressed from G1 to mid-M arrest and assessed for Rad52-GFP foci. As a control, a second culture was subjected to the same regime without auxin.

**Figure 4.**
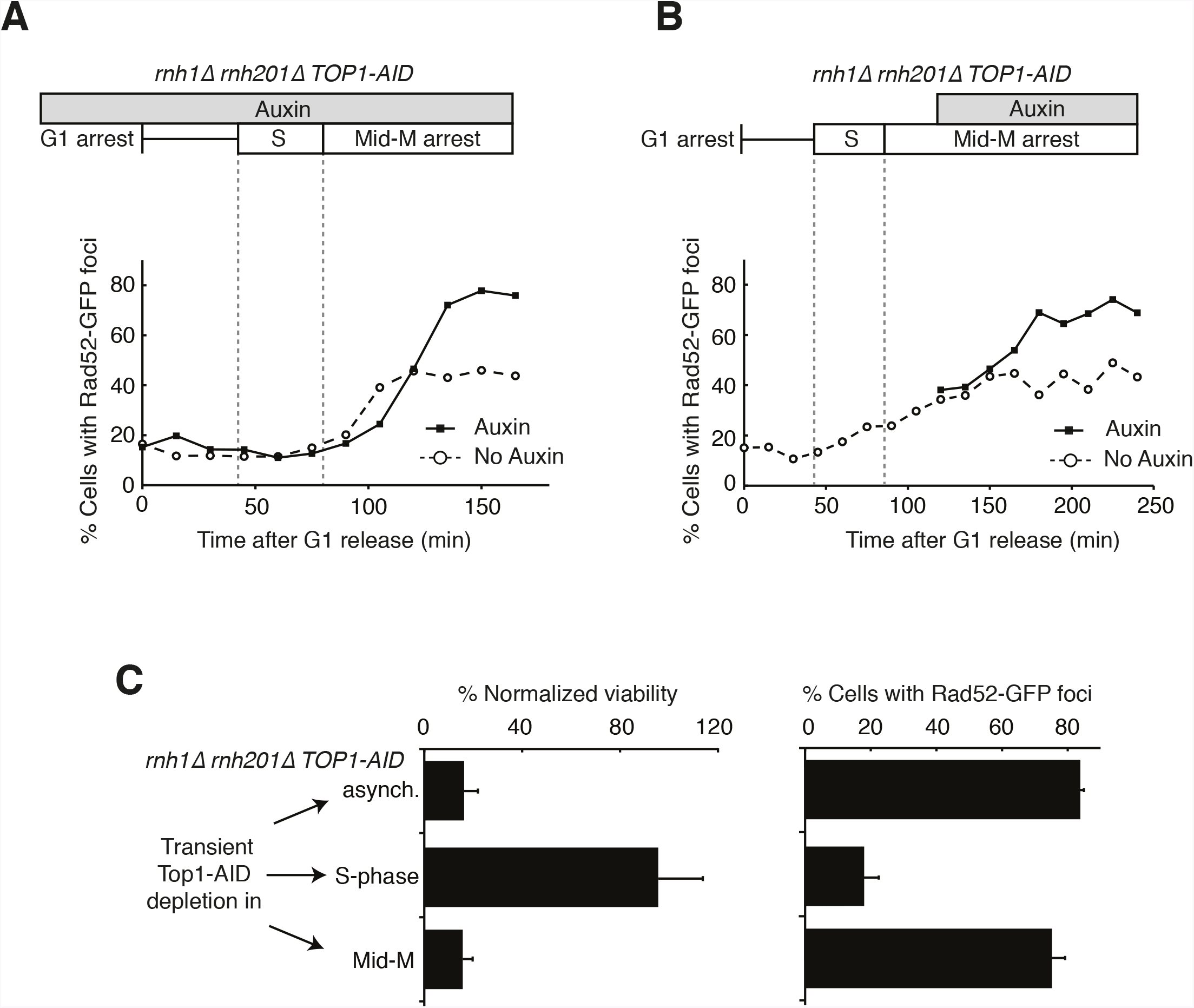
Depleting topoisomerase I causes lethal DNA damage in G2-M. (A) Depleting Top1 in rnh1 Δ rnh201Δ cells shows similar onset of Rad52-GFP at the S/G2-M border. Cultures of rnh1Δ rnh201Δ TOP1-AID cells were arrested in G1 using alpha factor, treated with auxin for 2 hours, then released into media containing nocodazole and auxin. Samples were taken at 15-minute intervals and 300 cells per time point were scored for Rad52-GFP foci. (B) Depleting Top1 in rnh1Δ rnh201Δ cells after completion of S-phase causes accumulation of Rad52-GFP foci. Cultures of rnh1Δ rnh201Δ TOP1-AID cells were released from alpha factor into nocodazole. Once cells had completed S-phase, auxin was added. Samples were taken at 30-minute intervals and 300 cells per time point were scored for Rad52-GFP foci. (C) Cultures of rnh1Δ rnh201Δ TOP1-AID cells were allowed to divide asynchronously, arrested in S-phase using hydroxyurea, or arrested in Mid-M phase using nocodazole. Once cells were arrested, auxin was added for four hours. Left – Cells were then washed and plated on YPD for recovery. Viability was measured by normalizing colony-forming units from auxin-treated cells to untreated cells. Bars represent mean +/- standard deviation (*n=4*). Right – Cells are scored for Rad52-GFP foci. Bars represent mean +/- standard deviation (*n=3*, 300 cells scored per replicate)

The enhanced foci in the triple mutant exhibited the same kinetics of accumulation as the double mutant (Figure 4A). The fraction of triple mutant cells with Rad52-GFP foci in both cultures remained around 15 to 20 percent until the end of bulk S-phase, similar to the *rnh1*Δ *rnh201*Δ double mutant. At subsequent time points, the auxin-free triple mutant culture mimicked the double mutant, as foci rose to about 45 percent in G2-M. The fraction of Rad52-GFP foci in the triple mutant cells treated with auxin also rose in G2-M but to a higher value of about 75 percent. Taken together, the triple mutant exhibited qualitatively similar but quantitatively greater cell cycle and DNA damage defects relative to the double mutant, indicating that Top1 depletion enhanced the DNA damage phenotype caused by loss of RNase H activity.

The ability to conditionally inactivate Top1-AID allowed us to address the role of Top1 activity in the cell-cycle dependent appearance of Rad52-GFP foci and the connection between focus formation and lethality. Cultures of *rnh1*Δ *rnh201*Δ *TOP1-AID* cells were arrested in mid-M with nocodazole and then treated with auxin. Top1-AID was depleted within 30 minutes of addition of auxin (Supplemental Figures 3B and 3C). The fraction of cells with Rad52-GFP foci climbed to about 70 percent (Figure 4B). This result suggested that Top1 activity was required after the completion of bulk S-phase to prevent focus formation.

To address whether Rad52-GFP foci induced by Top1 depletion were correlated with lethality, asynchronously dividing *rnh1*Δ *rnh201*Δ *TOP1-AID* cells were transiently treated with auxin for four hours, washed with fresh media, and then plated onto nonselective plates. The fraction of cells that survived, relative to an untreated control, was around 16 percent, similar to the percentage of cells that did not have Rad52 foci when given the same treatment (Figure 4C). This correlation suggested that the persistent foci in the *rnh1*Δ *rnh201*Δ *TOP1-AID* cells represented lethal DNA damage that arose from inactivation of Top1 activity in G2-M. To test this hypothesis further, we asked whether the appearance of foci was temporally correlated with inviability. The *rnh1*Δ *rnh201*Δ *TOP1-AID* cells were first arrested in S- or mid-M phase with hydroxyurea or nocodazole, respectively. The arrested cells were treated with auxin to deplete Top1-AID activity. These cells were plated for viability on media lacking auxin, allowing the restoration of Top1-AID activity (Figure 4C). Transiently depleting Top1 in S-phase, in which foci levels remain unchanged (18%), did not lead to loss of viability. However, transiently depleting Top1 in mid-M phase, which led to elevated foci levels (75%), also led to a dramatic increase in lethality. These results suggest that depletion of Top1 activity in G2-M in cells lacking the RNases H leads to irreparable DNA damage in the vast majority of cells.

### PIF1-E467G enables repair of R-loop mediated DNA damage

The lethality of *rnh1*Δ *rnh201*Δ cells upon Top1 inactivation provided a powerful genetic tool to interrogate R-loop induced DNA damage. Suppressor mutations that allowed a strain lacking both RNases H and Top1 to survive could either prevent damage from occurring or allow that damage to become reparable. These suppressor mutations could inform on the processes that convert R-loops to DNA damage or the mechanisms by which R-loop mediated damage is repaired. To isolate these suppressors, we generated independent cultures of an *rnh1*Δ *rnh201*Δ *top1*Δ strain that contained a plasmid carrying the *RNH1* and *URA3* genes. Plating these cultures on 5-fluoroorotic acid (5-FOA) selected for cells that had lost the plasmid and thus carried a suppressor mutation that allowed them to divide despite their *rnh1*Δ *rnh201*Δ *top1*Δ genotype (Figure 5A).

**Figure 5.**
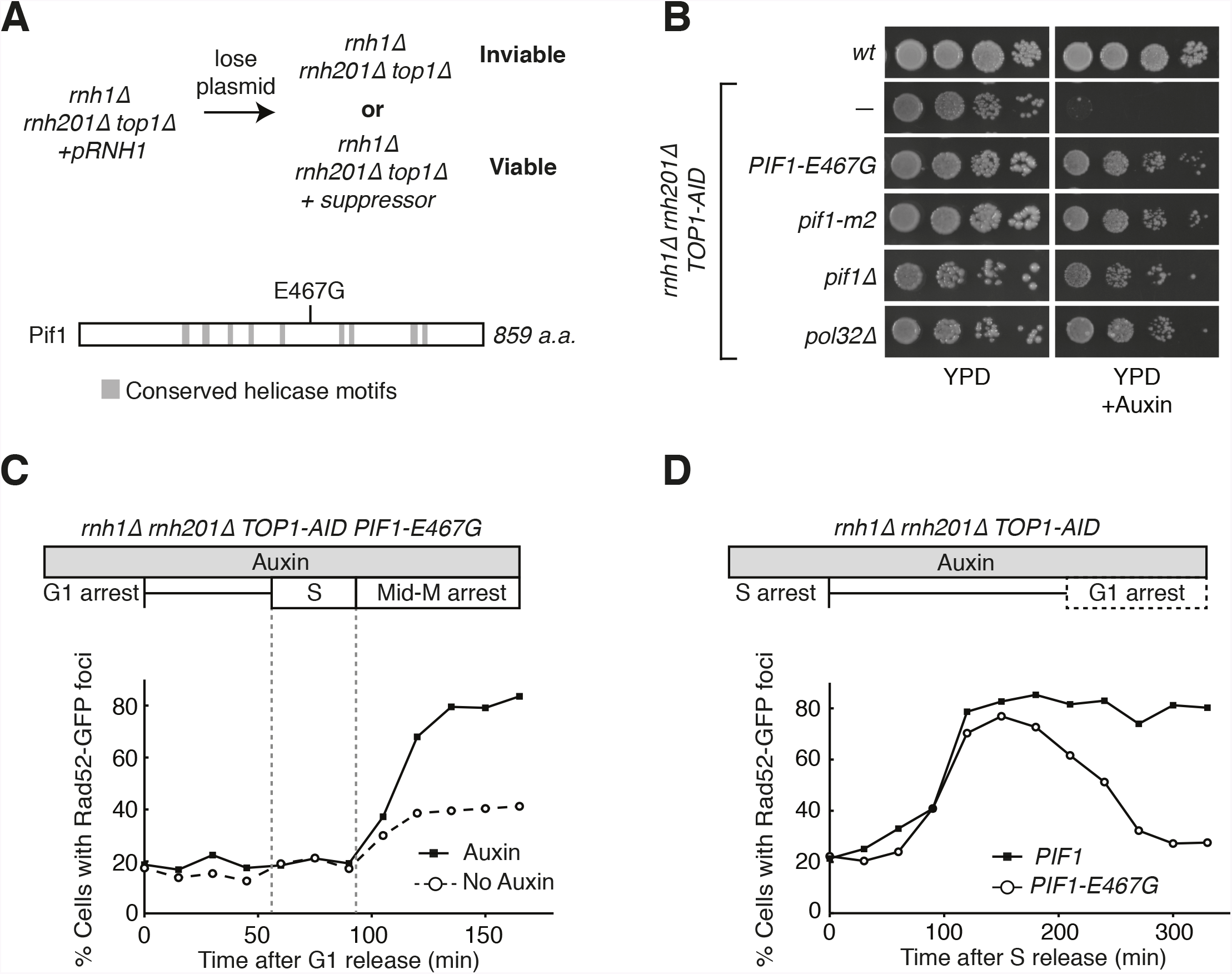
Pif1-E467G allows for repair of R-loop induced damage. (A) Top: schematic of genetic screen for suppressors of hybrid-induced lethality. Cells for the screen were rnh1Δ rnh201Δ top1Δ and carried a plasmid expressing RNH1 and URA3. Cultures were grown in non-selective media and plated onto 5-FOA to select for cells that had lost the plasmid and therefore gained suppressor mutations of rnh1Δ rnh201Δ top1Δ lethality. Bottom: schematic of Pif1 showing location of E467 relative to evolutionarily conserved SFI helicase motifs and motifs conserved between Pif1 and RecD, as previously published [24]. (B) Mutations in PIF1 and POL32 suppress auxin sensitivity of rnh1Δ rnh201Δ TOP1-AID cells. 10-fold serial dilutions of saturated cultures were plated onto YPD or YPD with auxin. (C) Pif1-E467G does not change accumulation of Rad52-GFP foci. Experiment in Figure 4A was repeated on rnh1Δ rnh201Δ TOP1-AID PIF1-E467G cells. (D) Pif1-E467G allows for repair of Rad52-GFP foci. Experiment in Figure 2B was repeated on rnh1Δ rnh201Δ TOP1-AID cells in the presence of auxin with or without PIF1-E467G.

DNA sequencing of the suppressor strains identified *PIF1-E467G.* E467G is a novel mutation in Pif1, a helicase with multiple roles in nuclear and mitochondrial DNA metabolism. This allele suppressed auxin sensitivity when introduced into *rnh1*Δ *rnh201*Δ *TOP1-AID* cells, indicating its responsibility for the suppression of lethality in *rnh1*Δ *rnh201*Δ *top1*Δ cells (Figure 5B). This allele was not found in any previously described domains of Pif1 (Figure 5A), prompting us to assess the ability of well-characterized recessive *PIF1* alleles to suppress the lethality of *rnh1*Δ *rnh201*Δ *top1*Δ genotype [24]. When introduced into *rnh1*Δ *rnh201*Δ *TOP1-AID*, both *pif1*Δ and *pif1-m2*, an allele that maintains mitochondrial but not nuclear functions of Pif1 [25], were able to suppress the auxin sensitivity of *rnh1*Δ *rnh201*Δ *TOP1-AID* cells. We conclude that *PIF1-E467G* likely inactivates a nuclear activity that is contributing to hybrid-induced lethality.

To determine whether *PIF1-E467G* prevented hybrid-induced DNA damage in *rnh1*Δ *rnh201*Δ *TOP1-AID* cells or allowed for its repair, we monitored the appearance and disappearance of Rad52-GFP foci in synchronously dividing cells. A culture of *rnh1*Δ *rnh201*Δ *TOP1-AID PIF1-E467G* cells was arrested in G1 (alpha factor) and treated with auxin to deplete Top1-AID. The culture was switched into media containing nocodazole and auxin to perpetuate Top1-AID depletion and allow progression through the cell cycle until arrest in mid-M phase (Figure 5C, Supplemental Figure 4A). The pattern of appearance of Rad52-GFP foci in this culture showed a strong similarity to the parent strain expressing wild-type Pif1, with no increase in foci until the completion of bulk S-phase. The fraction of cells containing Rad52-GFP foci then rose to 80 percent with auxin treatment and 40 percent without. These results suggest that the *PIF1-E467G* allele does not prevent hybrid induced DNA damage.

We next compared the appearance and disappearance of Rad52-GFP foci in *rnh1Δ rnh201Δ TOP1-AID* and *rnh1*Δ *rnh201*Δ *TOP1-AID PIF1-E467G* strains as they progressed between S phase to the subsequent G1. Cultures of these cells were arrested in hydroxyurea to induce S-phase arrest and then treated with auxin to induce Top1-AID depletion. The cultures were switched into media with auxin and alpha factor to perpetuate Top1-AID depletion and allow progression through mitosis to the next G1 (Figure 5B, Supplemental Figure 4B). In contrast to *rnh1*Δ *rnh201*Δ *TOP1-AID* cells, which maintained 85% Rad52-GFP foci and never proceeded through anaphase, cells with *PIF1-E467G* gradually resolved most of their foci as they completed mitosis. We therefore conclude that *PIF1-E467G* allows for the repair of hybrid-induced damage.

### Novel alleles of RNA polymerase I enable repair of R-loop mediated DNA damage

Our genetic screen also identified *RPA190-K1482T and RPA190-V1486F,* two novel alleles of Rpa190, the largest subunit of RNA pol I (Figure 6A). These residues map to the “jaw” domain of the RNA Pol I complex [26,27]. They are distinct from a previously tested Rpa190 allele that has been shown to suppress *rnh1*Δ *rnh201*Δ *top1*Δ inviability (*rpa190-3)*, and which is found closer to the dNTP entry pore of the polymerase [9,28]. *RPA190-K1482T* and *RPA190-V1486F* share the suppression phenotypes of *PIF1-E467G*. Both *RPA190* alleles suppress the auxin-induced inviability of *rnh1*Δ *rnh201*Δ *TOP1-AID* cells (Figure 6A). Similarly, neither allele prevents the accumulation of high levels of foci, but rather allow for their repair (Figures 6C and 6D, Supplemental Figures 5A and 5B). Given the specificity of RNA Pol I for transcribing regions of the rDNA locus, these results strongly suggest that the lethality in *rnh1*Δ *rnh201*Δ *top1*Δ is due to irreparable hybrid-induced DNA damage in the ribosomal repeats on chromosome XII and that altering the function of RNA polymerase I can allow this damage to be repaired.

**Figure 6.**
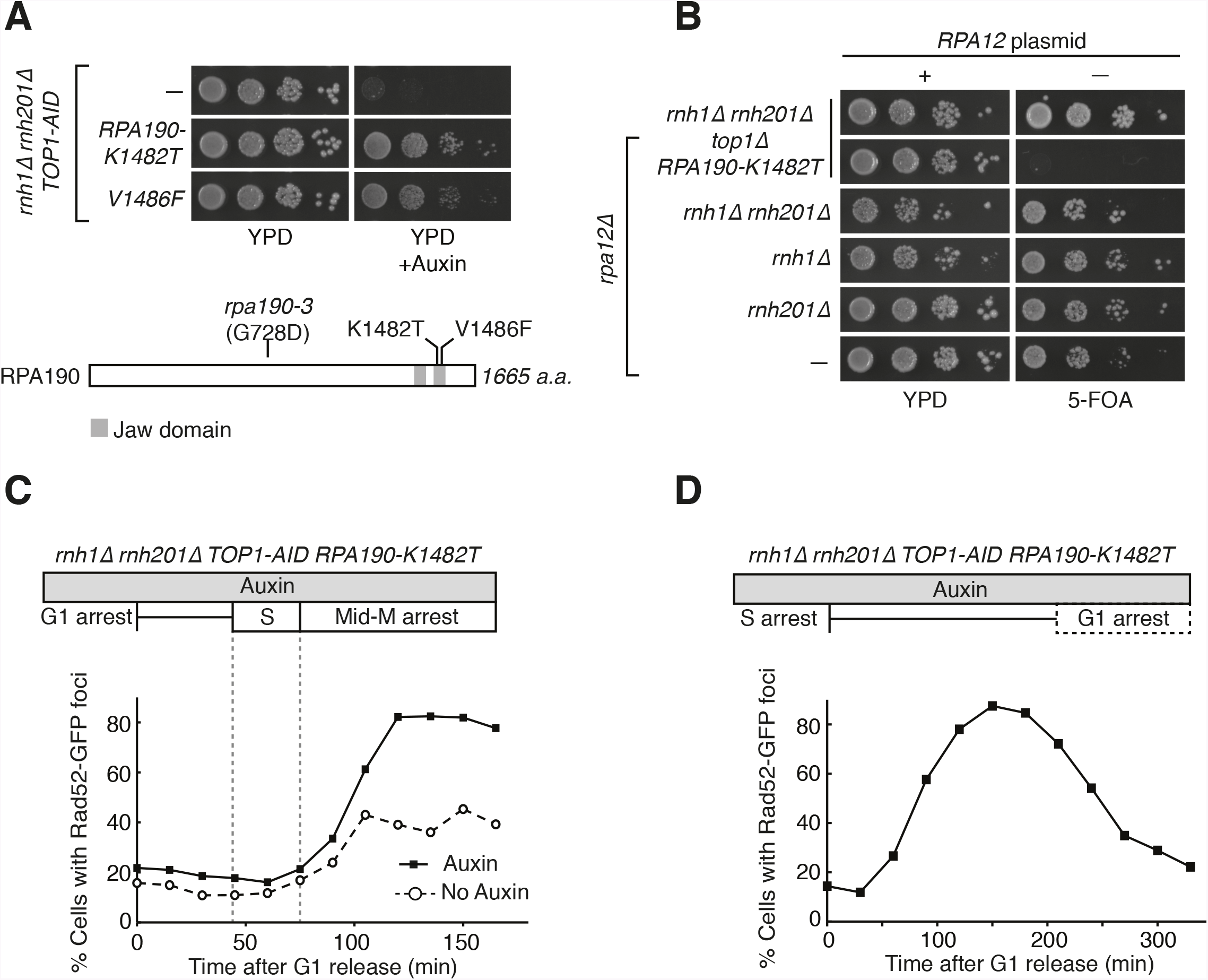
RPA190 mutants allow for repair of R-loop induced damage. (A) Top: Rpa190-K1482T and -V1486F suppress auxin sensitivity of rnh1Δ rnh201Δ TOP1-AID cells. 10-fold serial dilutions of saturated cultures were plated onto YPD or YPD with auxin. Bottom: Schematic of Rpa190 showing the location of mutations and the jaw domain, as previously published [26,27]. (B) Rpa12 is required in rnh1Δ rnh201Δ TOP1-AID RPA190-K1482T. Cells carrying a plasmid expressing RPA12 and URA3 were plated onto media lacking uracil (-URA, selects for plasmid) or media containing 5-floroorotic acid (5-FOA, selects for plasmid loss). 10-fold serial dilutions are shown. (C) Rpa190-K1482T does not change accumulation of Rad52-GFP foci. Experiment in Figure 4A was repeated on rnh1Δ rnh201Δ TOP1-AID RPA190-K1482T cells. (D) Rpa190-K1482T allows for repair of Rad52-GFP foci. Experi-ment in Figure 2B was repeated on rnh1Δ rnh201Δ TOP1-AID RPA190-K1482T cells in the presence of auxin.

To begin to understand the mechanism by which these alleles modify the activity of RNA pol I, we turned to previously reported crystal structures of the RNA pol I complex. Residues K1482 and V1486 in Rpa190 are found along potential contacts with the non-essential RNA pol I subunit Rpa12 (Supplemental Figures 6A-6C). Rpa12 has a role in promoting RNA Pol I backtracking and transcript termination [29,30]. This backtracking activity may affect genome stability – recent studies in bacteria have shown that backtracked RNA polymerases can cause R-loop dependent DSBs due to co-directional collisions with replisomes [31]. The proximity of Rpa12 to our suppressor mutations suggested that Rpa12 activities might be important for repair of the hybrid-induced damage. To test this idea, we knocked out Rpa12 in *rnh1*Δ *rnh201*Δ *top1*Δ *RPA190-K1482T* cells (Figure 6B). Deleting Rpa12 in these cells was lethal, indicating that our new *RPA190* alleles depend upon Rpa12, and by inference, polymerase backtracking or termination to repress hybrid-induced lethality.

### Break induced replication is responsible for inability to repair damage

Taken together, our studies implicate Pif1, the rDNA locus, and a G2-M specific process in the cell’s inability to repair hybrid-induced damage. These observations led us to question the role of break-induced replication (BIR) in hybrid-mediated instability. BIR is an HR-dependent repair process that occurs in G2-M when only one end of a DSB is available for recombination [32]. This end can be captured by homologous sequences and used as a primer for replication [33]. Break-induced replication is extremely processive and can extend the length of entire chromosomes, in part due to the contribution of the Pif1 helicase [34,35].

First, we asked how the *PIF1-E467G* allele functions in BIR. To do this, we introduced the allele into a previously characterized *in vivo* system for assessing BIR [36]. Briefly, these strains carry an inducible HO endonuclease cut site centromere-distal to an incomplete *URA3* gene on chromosome V. Upon induction of a DSB, a telomere needs to be added to the chromosome to restore viability to the cell. This repair can proceed with BIR using homology from an incomplete *URA3* repair template on the opposite arm. The repair template is situated in a position that requires either 30 or 80 kb of BIR for telomere addition, the latter being less efficient. Repair by BIR results in a fully functional *URA3* gene, thereby conferring the ability to grow on media lacking uracil [36].

In a wild-type strain, the frequency of BIR using the 30kb template is approximately 12%, consistent with previously published results (Figure 7A). In the absence of a functional Pif1 allele (*pif1-m2)*, repair by BIR drops to approximately 5%. Similarly, the frequency of BIR in cells carrying *PIF1-E467G* dropped to around 3%. In the 80kb repair template strain, a similar pattern in BIR efficiency was seen (Figure 7B). Strains carrying the *pif1-m2* or *PIF1-E467G* alleles saw drops in BIR frequency from approximately 5% to 1%. This led us to conclude that *PIF1-E467G* inhibits BIR. We observed no change in repair events in *top1Δ* cells, indicating that the absence of Top1 does not interfere with BIR.

**Figure 7.**
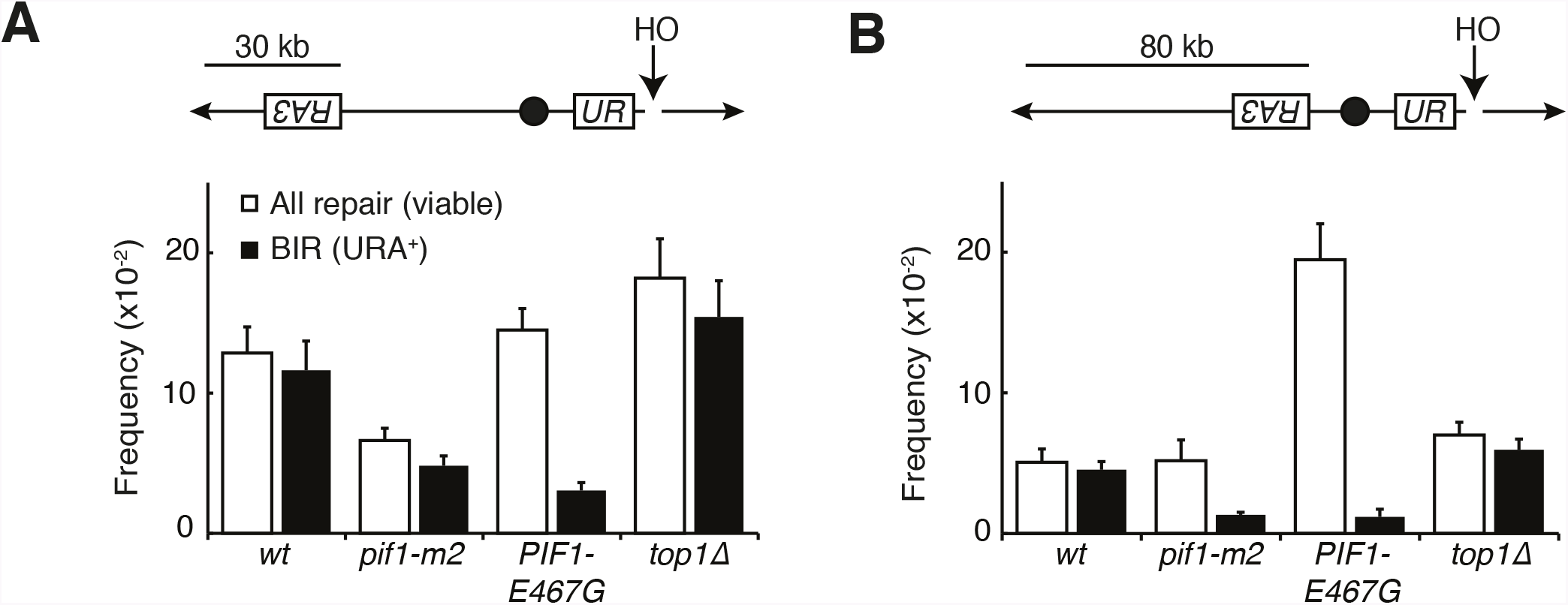
Pif1 mutants inhibit break-induced replication. (A) Top: Schematic of 30kb repair tem-plate strain. The HO endonuclease is under control of a GAL promoter. In the presence of galactose, it is expressed, inducing a DSB on chromosome V. Sequences telomeric to the HO cut site are non-essential. Homology between the two incomplete URA3 fragments allows for BIR and subsequent telomere addition. Bottom: Frequencies of repair. The percentage of cells that are viable on galactose (compared to total cells plated on non-DSB inducing YPD) indicates the frequency of all repair events. The subset of those cells that grow on media lacking uracil (URA+) indicates the frequency of BIR events. (B) As in (A), but with a repair template 80kb from the telomere. Bars represent mean +/- standard deviation (*n=4*).

The common phenotype for *PIF-E467G* and *pif1-m2* was inhibition of BIR, suggesting that this inhibition was responsible for allowing repair of the otherwise lethal hybrid-induced DNA damage. To test this model further, we deleted *POL32* in *rnh1*Δ *rnh201*Δ *TOP1-AID* cells (Figure 5B). Pol32 is a non-essential subunit of the primary BIR polymerase (Polδ), and is required for replication fork processivity [35]. These strains were no longer sensitive to auxin, corroborating the conclusion that inhibiting BIR is sufficient to allow for repair of otherwise lethal hybrid-induced DNA damage. Taken together, these results suggest that the attempt to repair hybrid-induced damage in the rDNA by BIR leads to an irreparable state.

*PIF1-E467G* promotes an alternative pathway for repair. In wild-type cells, almost all repair events after DSB induction used BIR. In contrast, *PIF1-E467G* cells had a near wild-type level of viability after DSB induction but used BIR only 20 percent of the time (Figures 7A and 7B). Thus, BIR was compromised, but an alternative pathway stepped in to promote telomere addition. As expected if this repair was independent of BIR, this alternative pathway was as efficient with the 80kb repair template as it was in cells with the 30 kb repair template; *PIF1-E467G* cells in the 80 kb repair strain had a total level of repair that was 5-fold greater than wild-type cells but only six percent of which was repaired using BIR. This alternative pathway was poorly activated in *pif1-m2* cells, as evidenced by a depressed level of viability. The increase in repair was not due to increased non-homologous end joining, as less than one percent of cells of all genotypes retained a telomeric drug resistance marker (Supplemental Figures 7A and 7B). Additionally, all strains efficiently induced DSBs, since PCR primers surrounding the cut site failed to amplify DNA after HO induction (Supplemental Figure 7C). Further analyses will be necessary to identify the alternative pathway for repair of hybrid-induced damage.

## Discussion

Our observations in this study suggest that the absence of the two RNases H leads to DNA damage that is difficult to repair after completion of S-phase. First, we observe an increase in Rad52 foci only in cells lacking both RNases H (*rnh1*Δ *rnh201*Δ). The increase in foci begins at the exit from S-phase and continues until mid-M. We show that this damage induces a significant pre-anaphase delay by activating the Rad9-dependent checkpoint. When measured in a bulk population, foci disappear slowly such that even after most cells have lost foci and completed cell division, a subset of cells retain foci and remain arrested pre-anaphase. This phenotype is indicative of an inability to efficiently repair damage. The depletion of Top1 in *rnh1*Δ *rnh201*Δ cells appears to exacerbate this problem by generating more foci that lead to lethal, irreparable damage and permanent arrest. Importantly, disrupting factors that modulate BIR allows for repair of these foci and restores cell division without reducing the initial level of damage. This demonstrates that the inability to repair damage, not the level of damage *per se*, in *rnh1*Δ *rnh201*Δ cells is the root cause of the inability to proceed through the cell cycle. Taken together, these results suggest that DNA:RNA hybrids inhibit DNA repair and that a critical role of the RNases H is to remove hybrids so that efficient repair can occur.

While hybrids have been recognized for many years as agents of genome instability, most studies have focused on their ability to generate DNA damage rather than their ability to alter DNA repair. However, a number of observations support our hypothesis of R-loops as inhibitors of repair. Inactivation of RNase H2 (*rnh201*Δ) by itself leads to large increases of genomic hybrids and loss of heterozygosity compared to wild-type [3,5]. This result suggests that loss of RNase H2 generates elevated levels of DNA damage. However, dramatically elevated Rad52 foci are only observed in *rnh1*Δ *rnh201*Δ cells – not *rnh201*Δ cells. We suggest that damage may be repaired rapidly in cells lacking RNase H2, while damage is repaired slowly in cells lacking both RNases H, causing foci to accumulate and persist.

Additionally, *sin3*Δ cells have elevated R-loops and hybrid-mediated genome instability. Inactivation of RNase H1 in *sin3*Δ cells increases genome instability further, but skews the events from chromosome repair to chromosome loss [4]. This result also supports a role for RNase H1 as critical in allowing repair of hybrid-induced damage. Presumably, under conditions of elevated hybrid formation such as *sin3*Δ, inactivation of RNase H1 alone is sufficient to cause a repair problem.

An insight into a potential role for the RNases H in DNA repair comes from a key observation in this study: lethality in *rnh1*Δ *rnh201*Δ cells when they are depleted of Top1 can be suppressed by mutations in Pif1 or Pol32 that inhibit BIR. Given the severity of the lethality – 88% cell death in a single cell cycle – this result suggests that BIR is a major pathway for the repair of R-loop induced damage in RNase H deficient cells. Consistent with this conclusion, previous studies mapped recombination events genome-wide in RNase H single and double mutants and found that 50% of the repair events occurred through BIR [5]. Furthermore, the percent of repair by BIR was elevated about 5 fold in the double mutant compared to either of the singles or wild type, although validation of this difference awaits a larger sample size.

To explain the BIR bias, we suggest that the RNases H remove hybrids from chromosomes both before and after R-loops induce DSBs (Figure 8A). Conversely, in the absence of RNase H activity, more DSBs are induced and hybrids persist at these DSBs. While one free end of the DSB may be properly processed by HR machinery, the presence of a hybrid on the opposite free end may block resection and/or invasion of homologous sequences. Ultimately, failure to capture the second free end of the DSB leads to BIR.

**Figure 8.**
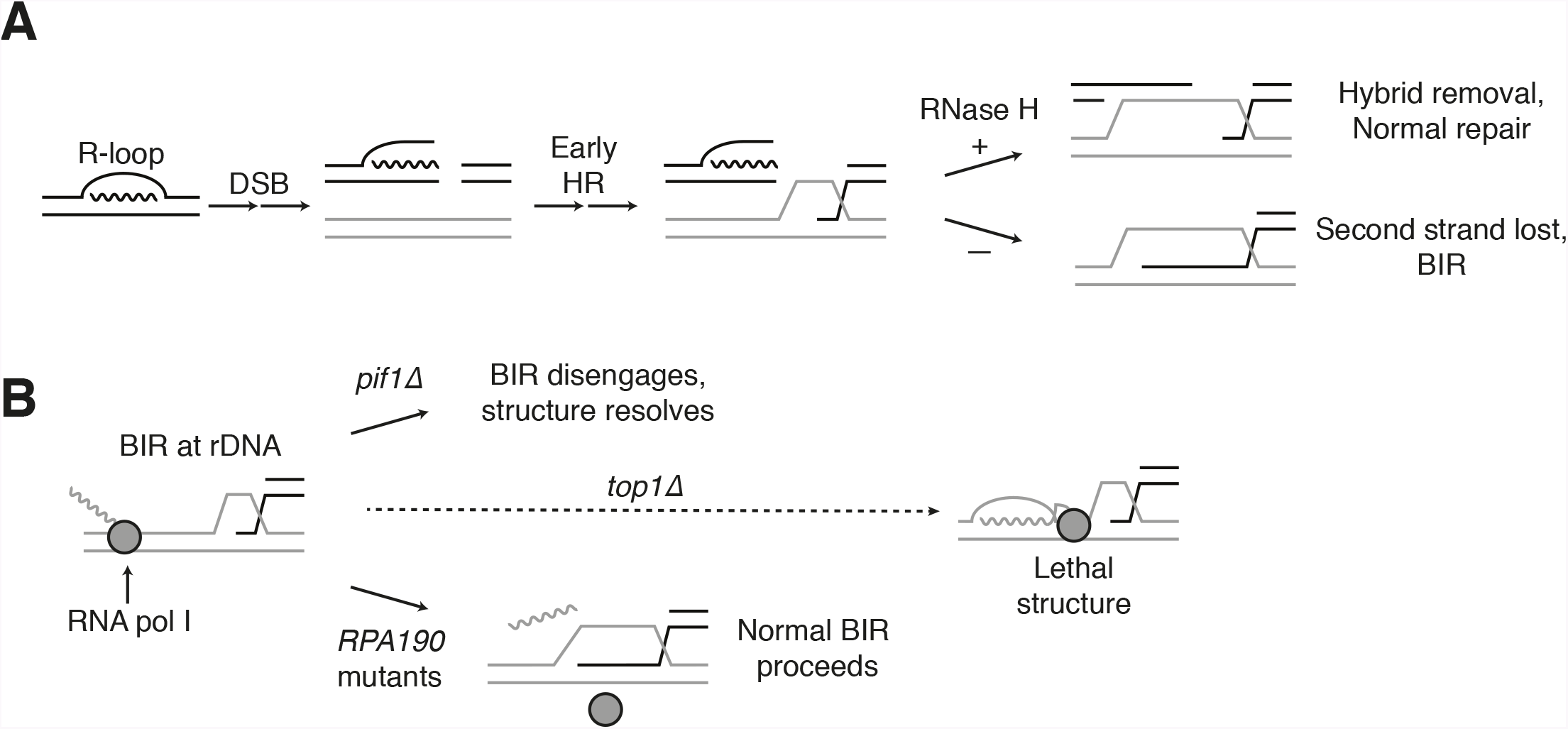
Proposed model for R-loop induced instability. (A) R-loops that cause DSBs persist at the break site. Early HR events (resection, homology search, strand capture and invasion) proceed as normal for one side of the break, but are inhibited by the presence of an R-loop on the opposite side. If RNase H1 or H2 act to clear the hybrid, repair can proceed as normal. If the hybrid persists, the second strand cannot be captured and the cell has no choice but to engage BIR. (B) BIR at the rDNA encounters replication blocks and slows. These replication blocks (over/under-winding, transcribing or stalled RNA polI, R-loops) are exacerbated in the absence of Top1, creating unresolvable structures that lead to cell death. PIF1 and POL32 mutants make BIR less processive, allowing BIR machinery to disengage before lethality occurs. The repair mechanism used after BIR is disengaged is unknown. RPA190 mutants allow for resolution of these structures, perhaps by disengaging RNA pol I through termination or backtracking activities.

Why does hybrid-induced BIR lead to cell-cycle arrest and a complete abrogation of repair when Top1 is depleted? An important clue comes from the fact that the lethality of *rnh1*Δ *rnh201*Δ *top1*Δ cells can be suppressed by mutations in RNA pol I, an enzyme whose function is limited to transcribing ribosomal DNA. The rDNA is the biggest source of R-loops in cells, accounting for almost 50% of all hybrids in yeast [3]. These results suggest that BIR at the rDNA may be particularly challenging in the presence of hybrids and even more challenging in the absence of Top1. We therefore propose that the induction of hybrids in the rDNA generates a barrier to the processivity of DNA replication during BIR, thereby slowing repair (Figure 8B). Further inactivation of Top1 causes elevated hybrids and stalled polymerases, which we hypothesize terminally block BIR replication fork progression. This trapped BIR intermediate is an aberrant structure that then leads to lethality. Interestingly, stalled replication forks have been observed at the rDNA locus in *rnh1*Δ *rnh201*Δ cells depleted of Top1 [9]. While these forks have been interpreted as being induced by aberrant DNA replication priming by the RNA moiety of R-loops, they equally well could have arisen from stalled BIR intermediates.

The mutations in RNA pol I that suppress the lethality of *rnh1*Δ *rnh201*Δ *top1*Δ map to the interface between subunits Rpa190 and Rpa12. This suppression is dependent upon Rpa12, a factor known to alleviate stalled polymerases either by promoting backtracking or transcription termination. Stalled RNA polymerases have been linked to R-loop dependent replication fork collisions that cause DSBs [31]. We therefore suggest that the *RPA190* suppressor mutations activate Rpa12, allowing it to remove stalled polymerases and possibly the associated hybrids, thereby removing the impediment for BIR imposed by Top1 (Figure 8B).

Our model for stalled BIR intermediates as the cause of lethality in *rnh1*Δ *rnh201*Δ *top1*Δ cells is supported by the molecular functions of Pif1 and Pol32. BIR is known to be a multi-step process in which strand invasion happens rapidly followed by a long delay before replication initiates [37]. This delay would explain the slow disappearance of hybrid-induced Rad52-GFP foci in *rnh1*Δ *rnh201*Δ cells. After this pause, Pif1 and Pol32 are required for processivity of the BIR replication fork. We suggest that inhibition of these two factors causes the invading strand to dissociate before it can reach the blocks imposed by the hybrids and/or RNA polymerases. A slower, alternative pathway, the identity of which remains unclear, can then repair the released strand.

In summary, we show that the RNases H play a critical role in promoting proper repair of hybrid-induced DNA damage, particularly in highly transcribed repetitive DNA. This conclusion came from directly limiting RNase H activity in cells and observing hybrid-induced BIR. Other conditions may also effectively limit RNase H activity. For example, many RNA biogenesis mutants induce hybrid formation and elevate genome instability [4]. The genome instability of these mutants can be suppressed by overexpression of RNase H, implying that RNase H activity becomes limiting when hybrid levels exceed a threshold. It has been assumed that this instability results only from increased damage induced by the persistence of R-loops. However, in light of our results, it is likely that limiting RNase H activity in these mutants also allows hybrids to promote BIR-induced genome instability. Finally, the particular sensitivity of the highly transcribed rDNA repeats is intriguing given that many cancer cells contain highly R-loop induced DNA damage, but also sites of improper repair.

## Materials and Methods

### Yeast strains, media, and reagents

Details on strain genotypes can be found in Supplemental Table 1. Plasmids used in this study can be found in Supplemental Table 2. YPD and synthetic complete minimal media were prepared as previously described [38]. For all cultures and plates using auxin, a one molar stock of 3-indoleacetic acid (Sigma-Aldrich) in DMSO was made and added to a final concentration of 500 µM. All auxin-treated experiments were compared to experiments that were mock-treated with equivalent volumes of DMSO. 5-fluorooritc acid (US Biological) was used at a final concentration of 1 mg/ml (w/v).

### Dilution plating assays

Cells were grown to saturation at 30^°^C in YPD. Cultures were then plated in 10-fold serial dilutions. Plates were incubated at 23^°^C. Representative images of experiments performed in duplicate or triplicate are shown.

### Chromosome spreads

Cells were collected and spheroplasted (0.1 M potassium phosphate [pH 7.4], 1.2 M sorbitol, 0.5 mM MgCl_2_, 20 mM DTT, 1.3 mg/ml zymolyase) at 37°C for 10 minutes or until >95% of cells lysed upon contact with 1% SDS. Spheroplasting reaction was stopped by washing and resuspension in a solution containing 0.1 M MES, 1 mM EDTA, 0.5 mM MgCl_2_, and 1M sorbitol (pH 6.4). Spheroplasts were placed onto slides and simultaneously lysed (1% Lipsol [v/v]) and fixed (4% paraformaldehyde [w/v], 3.4% sucrose [w/v]) by spreading solutions together across the slides using a glass pipette. then performed as previously described [7].

### Synchronous releases

Cells were grown to mid-log phase at 23^°^C in YPD. For G1 releases, alpha factor (Sigma-Aldrich) was added to 10^−8^ M and cultures were incubated for approximately 3.5 hours, until >95% of cells were visually confirmed to be arrested in G1. Cultures were split and treated with auxin or mock treated for two hours. Cells were then collected and washed six times in 1mL of YPD containing 0.1 mg/ml Pronase E (Sigma-Aldrich), with or without auxin depending on treatment. Cells were then resuspended in YPD containing nocodazole (Sigma-Aldrich) at a concentration of 15 µg/ml, with or without auxin, depending on treatment. Cultures were then grown at 23^°^C.

For S-phase releases, cells were arrested in hydroxyurea (Sigma-Aldrich) at a final concentration of 200 mM for 3 hours at 23^°^C. Cultures were split and treated with auxin or mock treated for 1.5 hours. Cells were washed 6x 1mL with YPD and released into YPD containing 10^−8^ M alpha factor (with or without auxin, depending on treatment). Cultures were then grown at 23^°^C. Note that all strains used in time-courses were *bar1Δ* to allow for greater sensitivity to alpha factor.

### Flow cytometry

Fixed cells were washed twice in 50 mM sodium citrate (pH 7.2), then treated with RNase A (50 mM sodium citrate [pH 7.2]; 0.25 mg/ml RNase A; 1% Tween-20 [v/v]) overnight at 37^°^C. Proteinase K was then added to a final concentration of 0.2 mg/ml and samples were incubated at 50^°^C for 2 hours. Samples were sonicated for 30s or until cells were adequately disaggregated. SYBR Green DNA I dye (Life Technologies) was then added at 1:20,000 dilution and samples were run on a Guava easyCyte flow cytometer (Millipore). 20,000 events were captured for each time point. Quantification was performed using FlowJo analysis software.

### Microscopy

Asynchronously and synchronously dividing cells were collected and resuspended in fixative (paraformaldehyde 4% [w/v] and sucrose 3.4% [w/v]) for 15 minutes at room temperature followed by washing and storage in 0.1 M potassium phosphate (pH 7.4), 1.2 M sorbitol. When indicated, nuclei were visualized by brief permiabilization of fixed cells with 1% Triton X 100 (v/v) followed by staining with DAPI at final concentration of 1 µg/ml. Scoring and image acquisition was with an Axioplan2 microscope (100× objective, numerical aperture [NA] 1.40; Zeiss, Thornwood, NY) equipped with a Quantix CCD camera (Photometrics, Tucson, AZ).

### Western blotting

Western blots were performed as previously described [39]. Primary antibodies used were a mouse monoclonal anti-V5 used at a 1:5000 dilution (Invitrogen) and a mouse monoclonal anti-Tub1 used at 1:20,000 dilution. Secondary antibody used was an HRP-conjugated goat anti-mouse at 1:20,000 (BioRad).

### Genetic Screen

Multiple independent cultures of strain JA271a were grown to saturation in YPD to allow for loss of plasmid pRS316-RNH1 (*RNH1 CEN URA3*). Cultures were diluted and plated so that dozens of colonies formed on each plate. Frequency of 5-FOA resistance was approximately 10^−7^. Colonies were then confirmed to be *rnh1*Δ *rnh201*Δ *top1*Δ, URA^−^, *grande*(able to grow on glycerol as the sole carbon source, indicating functional mitochondria), and not carry any temperature sensitivities. Genomic DNA was extracted and libraries were prepared using an Illumina TruSeq kit. Libraries were multiplexed and sequenced with 14-fold minimal coverage. Sequences were mapped to an S288c reference genome and SNPs were called relative to the parental JA271a strain. All three suppressors discussed here were built into *rnh1*Δ *rnh201*Δ *top1*Δ strains to confirm genetic linkage before being built into *TOP1-AID* strains.

### BIR assay

Experiments were performed as previously described [36]. Strains were grown on YPD +cloNAT plates. Individual colonies were picked and serially diluted in water so that ~200 cells were plated onto YPD and ~2000 cells were plated on YP-GAL. Cells that grew on YP-GAL were counted then replica-plated onto SC –URA and YPD +cloNAT plates. Total survivors were calculated by dividing the number of colonies that grew on YP-GAL by the number that grew on YPD, adjusting for 10-fold dilution. Similar calculations were performed on SC –URA and YPD +cloNAT plates to determine rates of BIR and NHEJ, respectively.

## Acknowledgments

We thank A. Zimmer, L. Costantino, H. Tapia, B. Robison, R. Lamothe, A. Muir, and S. Kim for comments on the manuscript. We would also like to thank V.A. Zakian for plasmids and J.E. Haber and for strains and helpful discussions.

---

**Supplemental Figure 1.**
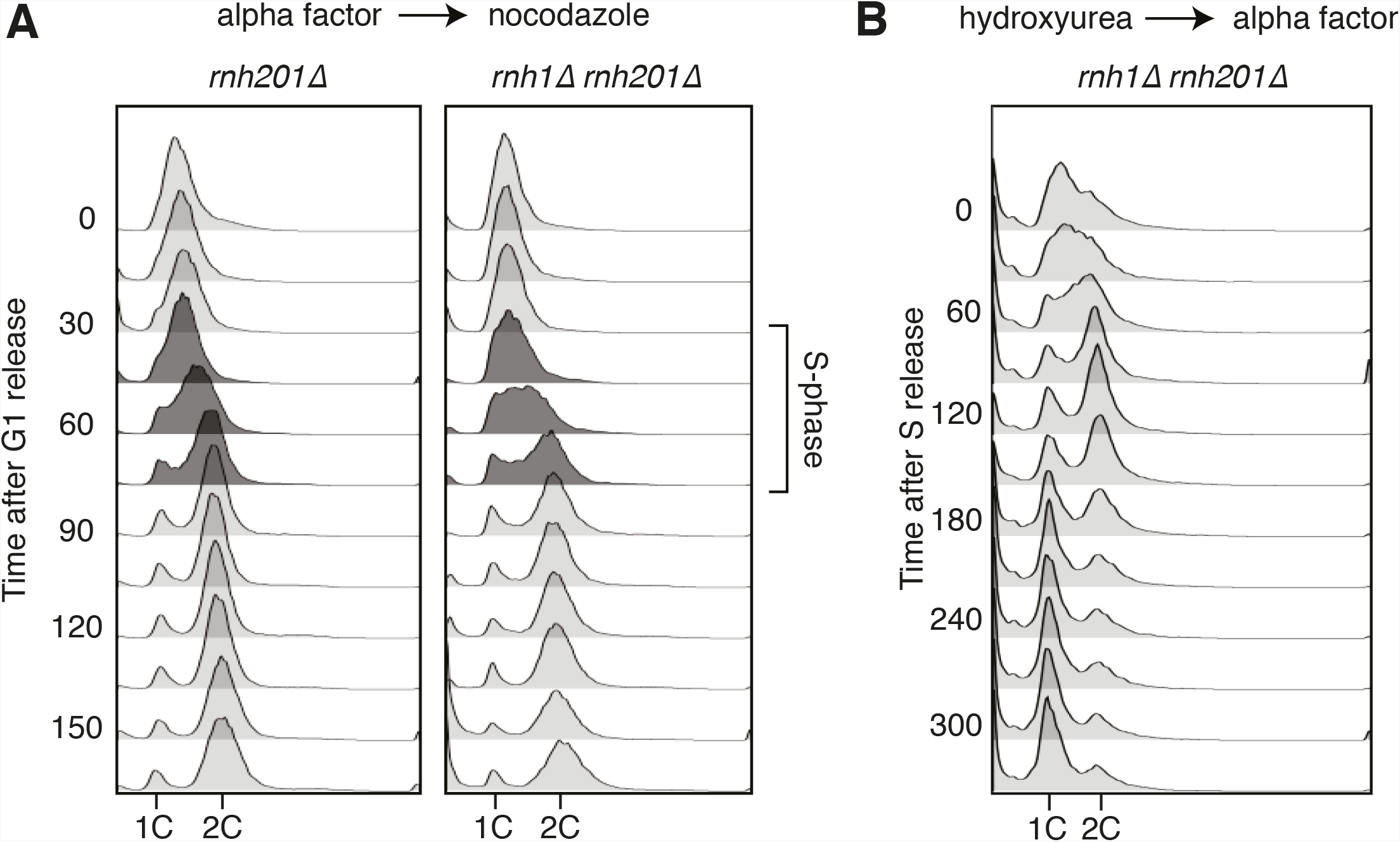
(A) Flow cytometry of rnh201Δ and rnh1Δ rnh201Δ cells released from alpha factor into nocodazole. Corresponds to figure 1C. (B) Flow cytometry of rnh1Δ rnh201Δ cells released from hydroxyurea into alpha factor. Corresponds to figure 2B.

**Supplemental Figure 2.**
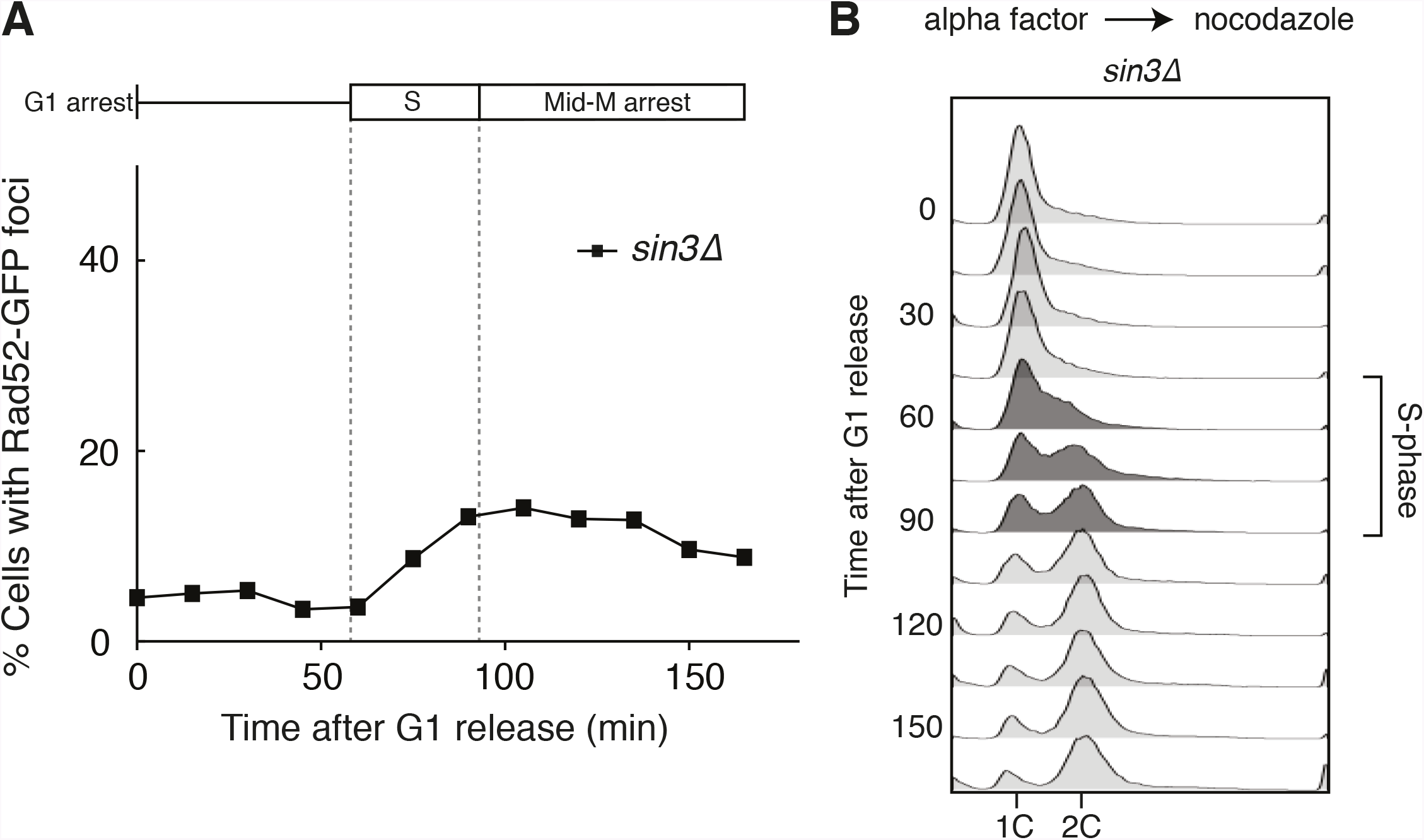
Deleting SIN3 causes increased foci in S phase. (A) Cultures of sin3 Δ cells were arrested in alpha factor and released into nocodazole. Samples were taken at 15 minute intervals, and cells were scored for Rad52-GFP foci. (B) Flow cytometry of cells in (A).

**Supplemental Figure 3.**
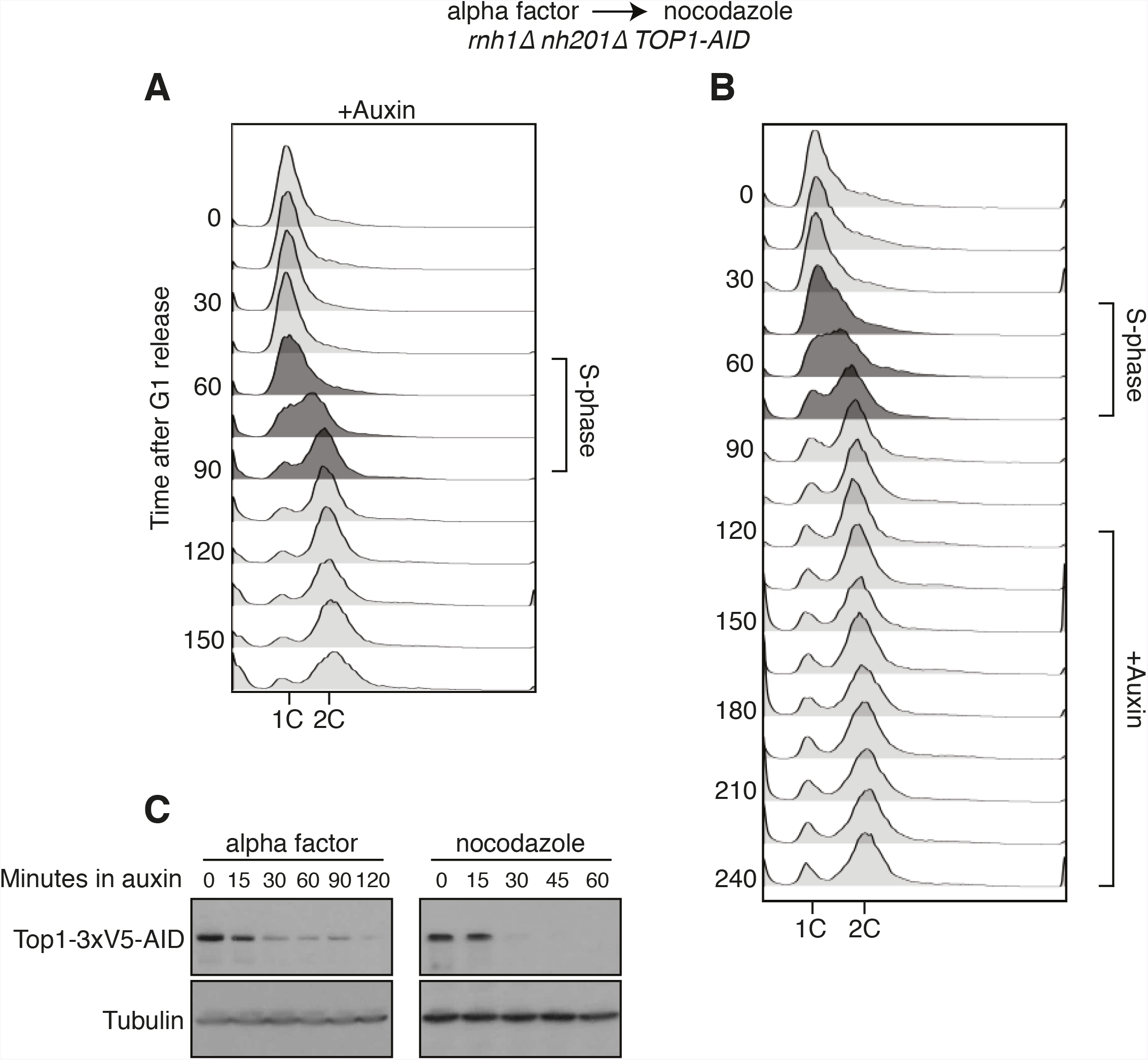
Details on rnh1Δ rnh201Δ TOP1-AID cells released from alpha factor into nocodazole. (A) Flow cytometry corresponds to figure 4A. Auxin is added to cells while they are arrested in alpha factor. Cells are then released into nocodazole and auxin, maintaining the state of Top1 depletion. (B) Flow cytometry corresponds to figure 4B. Auxin is added to cultures in nocodazole 120 minutes after release from alpha factor. (C) Cells arrested in alpha factor or nocodazole were treated with auxin. Samples were taken at the indicated time points and processed for western blotting. Top: Mouse anti-V5 antibody detects Top1-3xV5-AID. Addition of auxin depletes Top1 by two hours in alpha factor and 30 minutes in nocodazole. Bottom: Rabbit anti-tubulin antibody detects Tub1.

**Supplemental Figure 4.**
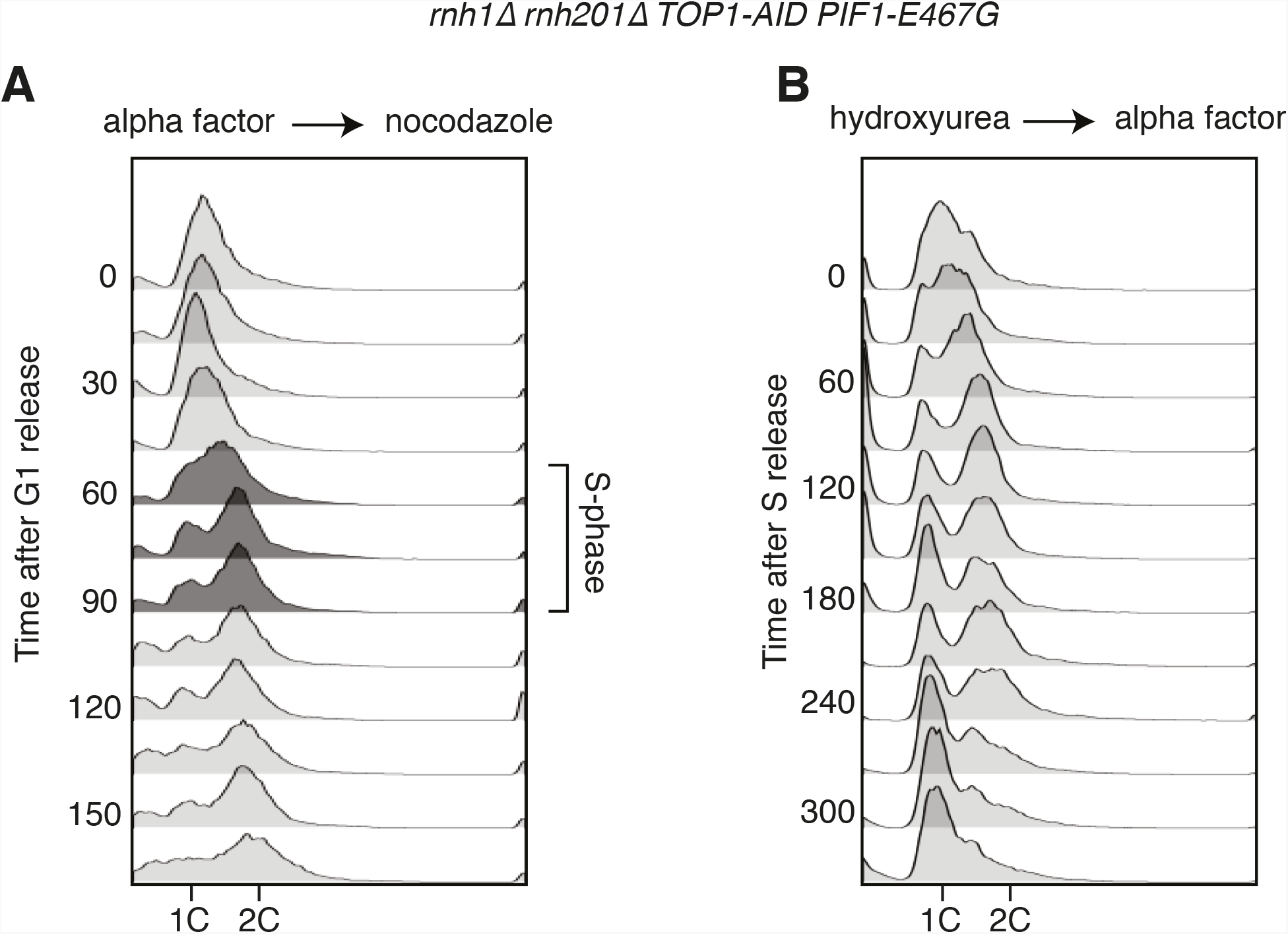
Flow cytometry on rnh1Δ rnh201Δ TOP1-AID PIF1-E467G cells. All cells have been treated with auxin for two hours before release and auxin is maintained in the culture after release. (A) Cells are released from alpha factor into nocodazole. Flow cytometry profiles correspond to figure 5C. (B) Cells are released from hydroxyurea into alpha factor. Corresponds to figure 5D.

**Supplemental Figure 5.**
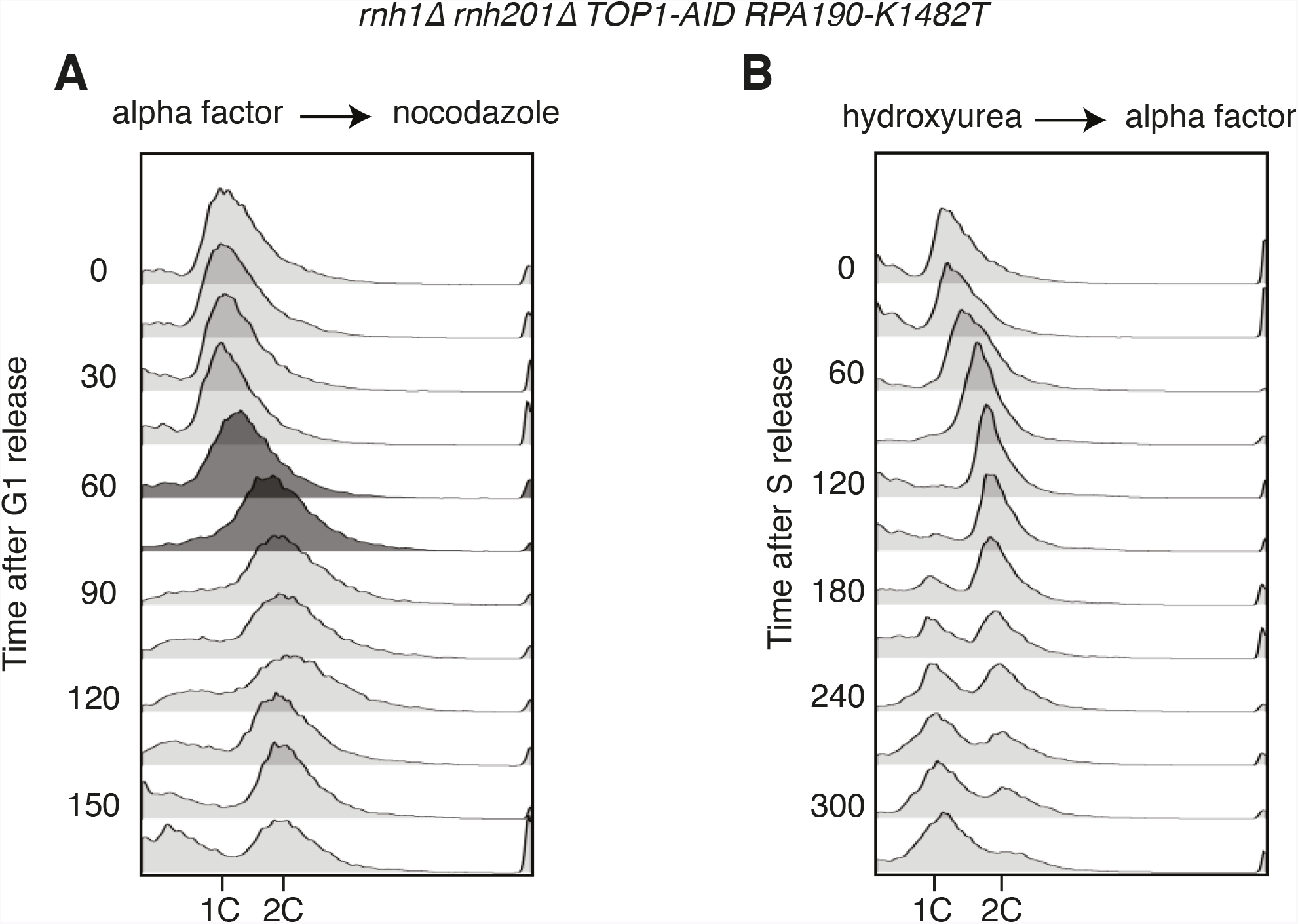
Flow cytometry on rnh1Δ rnh201Δ TOP1-AID RPA190-K1482T cells. All cells have been treated with auxin for two hours before release and auxin is maintained in the culture after release. (A) Cells are released from alpha factor into nocodazole. Flow cytometry profiles correspond to figure 6C. (B) Cells are released from hydroxyurea into alpha factor. Corresponds to figure 6D.

**Supplemental Figure 6.**
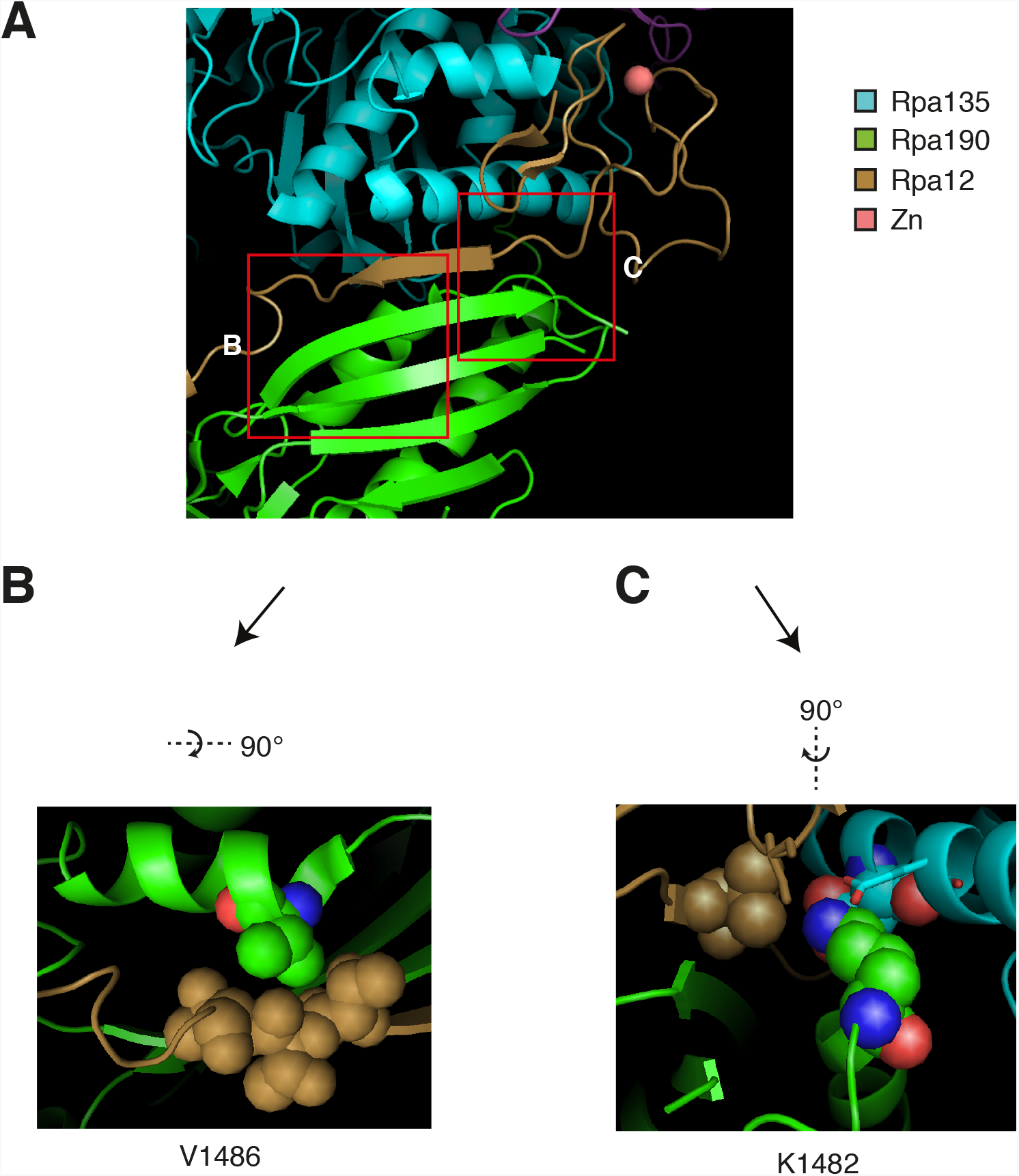
Structural analysis of Rpa190 in the context of the RNA pol I complex. Structure shown is from Protein Data Bank accession number 4C3I, as originally published in Fernández-Tornero, et al. (2013). (A) The “jaw” domain of Rpa190 (green) is shown with the N-terminal zinc-ribbon of Rpa12 (brown) and the “lobe” domain of Rpa135 (cyan). (B) The highlighted region rotated to see residue V1486 in its position on the alpha helix of RPA190 behind the beta sheet in the foreground of (A). Residue V1486 on Rpa190 is shown in spherical space along with residues T49, T50, and T51 of Rpa12. Steric clashes arise between these residues when Rpa190-V1486 is modified to phenylalanine. (C) The highlighted region rotated to see residue K1482 behind the beta sheet in the foreground of (A). Residue K1482 of Rpa190 is shown in spherical space along with Rpa135-D304 and Rpa12-V47 in the background. Also in close proximity are Rpa135-E307 and Rpa12-S6 shown as stick models in the foreground. Modifying Rpa190-K1482 to threonine increases the distance between these residues and possibly interrupts acid-base interactions anchored by the lysine residue.

**Supplemental Figure 7.**
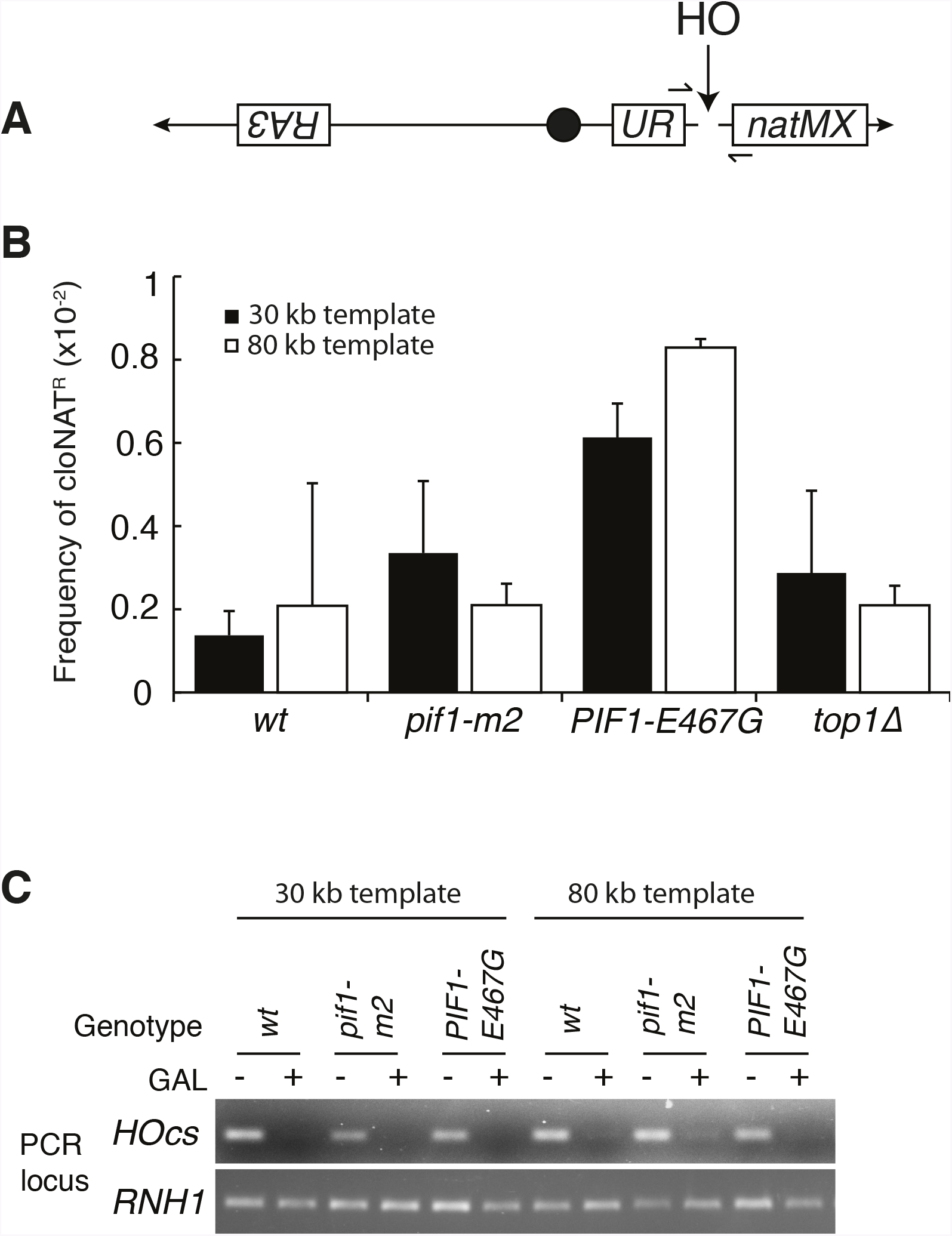
Mutants don’t affect non-homologous end joining or HO-induced DSBs. (A) More-detailed schematic of chromosome V constructs being used in Figure 7. A gene that confers resistance to the drug cloNAT (natMX) is placed telomeric to the HO cut site (HOcs). Primers designed to flank the HOcs are shown as well. (B) Frequencies of cloNAT resistance in various genotypes in both 30- and 80kb repair template strains. Viable colonies grown on galactose (Figure 7) were replica-plated to media containing cloNAT. Frequency is calculated by comparing cells that grew on cloNAT to total cells on YPD. Ability to grow on uracil and cloNAT resistance were mutually exclusive. Bars represent mean +/- standard deviation (n=4). (C) Saturated cultures of the indicated genotype were diluted into media containing dextrose or galactose and allowed to divide for six hours. Genomic DNA was extracted and PCR was performed at the HO cut site (HOcs) or the RNH1 locus as a positive control. PCR products were loaded onto an agarose gel and stained with ethidium bromide.

**Supplemental Table 1.** Strains used in this study

| Strain | Genotype | Reference |
| --- | --- | --- |
| JA30 | *MATa RAD52-GFP-HIS3* |  |
| JA373a | *MATa RAD52-GFP-HIS3 rnh1Δ::KAN* |  |
| JA320a | *MATa RAD52-GFP-HIS3 rnh201Δ::NAT* |  |
| JA338a | *MATa RAD52-GFP-HIS3 rnh1Δ::KAN rnh201Δ::NAT* |  |
| JA426b | *MATa RAD52-GFP-HIS3 rnh1Δ::KAN rnh201Δ::NAT rad9Δ::URA3* |  |
| JA382 | *MATa rad52Δ::LEU2 pRS316-RAD52* |  |
| JA378 | *MATa rad52Δ::LEU2 pRS316-RAD52 rnh1Δ::KAN* |  |
| JA380 | *MATa rad52Δ::LEU2 pRS316-RAD52 rnh201Δ::NAT* |  |
| JA376 | *MATa rad52Δ::LEU2 pRS316-RAD52 rnh1Δ::KAN rnh201Δ::NAT* |  |
| JA249a | *MATalpha rad9Δ::URA3* |  |
| JA255a | *MATa RAD52-GFP-LEU2 his3-Δ1::TIR1-HIS3 bar1Δ::hisG* |  |
| JA268b | *MATa RAD52-GFP-LEU2 his3-Δ1::TIR1-HIS3 bar1Δ::hisG sin3Δ::KAN* |  |
| JA238 | *MATa RAD52-GFP-LEU2 his3-Δ1::TIR1-HIS3 bar1Δ::hisG-URA3-hisG* |  |
| JA210 | *MATalpha RAD52-GFP-LEU2 his3-Δ1::TIR1-HIS3 bar1Δ::hisG-URA3-hisG*<br>*TOP1-3xV5-AID-KAN* |  |
| JA208 | *MATa RAD52-GFP-LEU2 his3-Δ1::TIR1-HIS3 bar1Δ::hisG-URA3-hisG*<br>*TOP1-3xV5-AID-KAN rnh1Δ::HYG* |  |
| JA209 | *MATa RAD52-GFP-LEU2 his3-Δ1::TIR1-HIS3 bar1Δ::hisG-URA3-hisG*<br>*TOP1-3xV5-AID-KAN rnh201Δ::NAT* |  |
| JA204a | *MATa RAD52-GFP-LEU2 his3-Δ1::TIR1-HIS3 bar1Δ::hisG-URA3-hisG*<br>*TOP1-3xV5-AID-KAN rnh1Δ::HYG rnh201Δ::NAT* |  |
| JA214 | *MATa RAD52-GFP-LEU2 his3-Δ1::TIR1-HIS3 bar1Δ::hisG-URA3-hisG*<br>*rnh201Δ::NAT* |  |
| JA207 | *MATa RAD52-GFP-LEU2 his3-Δ1::TIR1-HIS3 bar1Δ::hisG-URA3-hisG*<br>*rnh201Δ::NAT rnh1Δ::HYG* |  |
| JA205 | *MATa RAD52-GFP-LEU2 bar1Δ::hisG-URA3-hisG rnh1Δ::HYG rnh201Δ::NAT* |  |
| JA240 | *MATa RAD52-GFP-LEU2 his3-Δ1::TIR1-HIS3 bar1Δ::hisG*<br>*TOP1-3xV5-AID-KAN rnh1Δ::HYG rnh201Δ::NAT* |  |
| JA250a | same as JA240 except *rad9Δ::URA3* |  |
| JA302a | same as JA240 except *RPA190-K1482T* |  |
| JA303a | same as JA240 except *RPA190-V1486F* |  |
| JA304a | same as JA240 except *PIF1-E467G* |  |
| JA413a | same as JA240 except *pif1-m2* |  |
| JA421a | same as JA240 except *pif1Δ::URA3* |  |
| JA423a | same as JA240 except *pol32Δ::URA3* |  |
| JA271a | *MATa rnh1Δ::KAN rnh201Δ::NAT top1Δ::HYG RAD52-GFP-HIS3 pRS316-RNH1* |  |
| JA308a | *MATa rnh1Δ::HYG rnh201Δ::NAT top1Δ::HYG RAD52-GFP-HIS3 RPA190-K1482T* |  |
| JA309b | *MATa rnh1Δ::HYG rnh201Δ::NAT top1Δ::HYG RAD52-GFP-HIS3 RPA190-V1486F* |  |
| JA310 | *MATa rnh1Δ::HYG rnh201Δ::NAT top1Δ::HYG RAD52-GFP-HIS3 RPA190-K1482T* |  |
| JA394 | *MATa pRS316-RPA12 RAD52-GFP-HIS3 rpa12Δ::LEU2* |  |
| JA392a | *MATa pRS316-RPA12 RAD52-GFP-HIS3 rpa12Δ::LEU2 rnh1Δ::KAN* |  |
| JA393a | *MATa pRS316-RPA12 RAD52-GFP-HIS3 rpa12Δ::LEU2 rnh201Δ::NAT* |  |
| JA390 | *MATa pRS316-RPA12 RAD52-GFP-HIS3 rpa12Δ::LEU2 rnh1Δ::KAN rnh201Δ::NAT* |  |
| JA344a | *MATa pRS316-RPA12 RAD52-GFP-HIS3 rpa12Δ::LEU2 rnh1Δ::KAN rnh201Δ::NAT*<br>*top1Δ::HYG RPA190-K1482T* |  |
| JA325a | *MATa pRS316-RPA12 RAD52-GFP-HIS3 rnh1Δ::KAN rnh201Δ::NAT top1Δ::HYG*<br>*RPA190-K1482T* |  |
| yRA52 | *MATa::ΔHOcs::hisG ura3Δ851 trp1Δ63 leu2Δ::KAN hmlΔ::hisG HMR::ADE3*<br>*ade3::GAL::HO can1Δ::UR::HOcs::NAT, RA3::TRP1 (30kb)* | Anand, et al. 2014 |
| JA415a | same as yRA52 except *pif1-m2* |  |
| JA416a | same as yRA52 except *PIF1-E467G* |  |
| JA447a | same as yRA52 except *top1Δ::HYG* |  |
| yRA107 | *MATa::ΔHOcs::hisG ura3Δ851 trp1Δ63 leu2Δ::KAN hmlΔ::hisG HMR::ADE3 ade3::GAL::HO can1Δ::UR::HOcs::NAT, RA3::TRP1 (80kb)* | Anand, et al. 2014 |
| JA417a | same as yRA107 except *pif1-m2* |  |
| JA418a | same as yRA107 except *PIF1-E467G* |  |
| JA448a | same as yRA107 except *top1Δ::HYG* |  |

**Supplemental Table 2.** plasmids used in this study

| **Plasmid** | **Description** | **Reference** |
| --- | --- | --- |
| pRS316 | *CEN ARS URA3* low copy plasmid | Lab stock |
| pRS306 | *URA3* integrating lasmid | Lab stock |
| pRS316-*RAD52* | Rad52 complementation plasmid. | This study |
| pRS316-*RPA12* | Rpa12 complementation plasmid. | This study |
| pRS306-*RPA190-V1486F* | Integrating plasmid for creating *RPA190-K1482T*. | This study |
| pRS306-*RPA190-V1486F* | Integrating plasmid for creating *RPA190-V1486F*. | This study |
| pRS306-*PIF1-E467G* | Integrating plasmid for creating *PIF1-E467G*. | This study |
| pVS31 | Integrating plasmid for creating *pif1-m2* | Schulz and Zakian, 1994 |

